# Widespread naturally variable human exons aid genetic interpretation

**DOI:** 10.1101/2024.09.09.612029

**Authors:** Hannah Jacobs, Bram L. Gorissen, Jeremy Guez, Masahiro Kanai, Hilary K. Finucane, Konrad J. Karczewski, Christopher B. Burge

**Affiliations:** MIT Department of Biology, Cambridge, Massachusetts 02139, USA; Program in Medical and Population Genetics, Broad Institute of MIT and Harvard, Cambridge, Massachusetts 02142, USA; Novo Nordisk Foundation Center for Genomic Mechanisms of Disease, Broad Institute of MIT and Harvard, Cambridge, Massachusetts 02142, USA; Analytic and Translational Genetics Unit, Massachusetts General Hospital, Boston, Massachusetts 02114, USA

## Abstract

Most mammalian genes undergo alternative splicing. The splicing of some exons has been acquired or lost in specific mammalian lineages, but differences in splicing within the human population are poorly characterized. Using GTEx tissue transcriptomes from 838 individuals, we identified 56,415 exons which are included in mRNAs in some individuals but entirely excluded from others, which we term “naturally variable exons” (NVEs). NVEs impact three quarters of protein-coding genes, occur at all population frequencies, and are often absent from reference annotations. NVEs are more abundant in genes depleted of genetic loss-of-function mutations and aid in the interpretation of causal genetic variants. Genetic variants modulate the splicing of many NVEs, and 5’UTR and coding-region NVEs are often associated with increased and decreased gene expression, respectively. Together, our findings characterize abundant splicing variation in the human population, with implications for a range of human genetic analyses.

## Introduction

Nearly all human genes undergo alternative splicing, in which distinct mRNAs derived from different combinations of exons and splice sites are produced from the same gene^1^. Small changes in mRNA sequence can exert large effects, including addition/deletion of protein domains^2,3^, or production of functional versus nonfunctional isoforms, and dysregulation of splicing contributes to human disease^4,5^. Understanding variation in alternative splicing is therefore important in understanding physiological and disease mechanisms.

Evolutionarily conserved alternatively spliced exons tend to preserve protein reading frame^6^, and often exhibit tissue-specific regulation^7^. Additionally, comparison across mammalian species has identified thousands of cases of complete loss or gain of the splicing of exons^8^, with newly evolved splicing events more likely to occur in 5’ UTRs, where they are often associated with increased gene expression^8^. Gain or loss of the splicing of an exon in the coding sequence (CDS) may also influence expression by changing the reading frame, often triggering nonsense-mediated mRNA decay (NMD)^9,10^. However, the evolution of alternative splicing within the human species remains poorly understood.

Most alternative exons are generally assumed to be spliced in all individuals, perhaps with different inclusion levels, a pattern which we call ‘canonical’ alternative splicing. Some of these splicing differences may be driven by genetic variation. Large-scale genetic association studies have associated many single nucleotide polymorphisms (SNPs) with changes in the usage of alternatively spliced exons, designated splicing quantitative trait loci (sQTLs)^11,12^. However, these variants typically have modest effect sizes, and their molecular mechanisms are often unclear. Some ultra-rare genetic variants can lead to inclusion of previously unseen, “cryptic” exons or splice sites^13^. Some recent studies suggest that many exons are missing from transcriptome reference annotations, occurring only in a subset of transcriptomic datasets^11,14,15^. The availability of tissue transcriptomes from around 1000 individuals in the Genotype-Tissue Expression (GTEx) project presents an opportunity to characterize human variation at the exon level, asking how often the presence/absence of individual exons or splice sites varies between the transcriptomes of different individuals.

One key challenge in the study of unannotated transcripts is assessing which reflect technical variation or other types of noise^16,17^, which are reproducible, and which have phenotypic effects^18^. A cross-species analysis has suggested that many unannotated splicing events in longer-lived organisms may represent “transcript drift”, and are often nonfunctional until they begin to splice in at higher rates^19^. However, another recent study reasoned that lowly used exons that trigger NMD broadly impact gene expression in human lymphoblastoid cell lines, and can help interpret complex trait loci^10^.

Here, we define “naturally variable exons” (NVEs) as exons whose splicing is variable within the human population (i.e., not present in every individual), focusing on alternative “skipped” or “cassette” exons and alternative splice sites, which extend or truncate exons. We introduce a statistical approach that estimates the population frequency with which NVEs are meaningfully included (at levels above 5%) in a human cohort, generating a quantitative atlas of splicing variation across tissues. Using data from GTEx, we find that NVEs impact most human genes, are sometimes associated with changes in gene expression, and can aid in the interpretation of non-coding genetic variants.

## Results

### Generating a catalog of human NVEs

In order to generate a catalog of high-quality NVEs, we obtained mapped RNA-seq reads from 14,000 GTEx samples representing 49 tissues from 838 individuals^11^. To perform quality control and ensure accurate estimates of NVEs in the dataset, we devised a straightforward Bayesian approach. The method de-noises estimates by sharing signals across samples and enables estimation of the uncertainty in exon usage of each NVE.

To illustrate the method, consider an NVE alternative splice site used in some individuals as an alternative to a canonical or “cognate” splice site, used in all people. The proportional use of the NVE within an individual – the “percent spliced in’’ (PSI or Ψ) – can be estimated from the proportion of RNA-seq reads that span the NVE and cognate exon junctions (EJs) (**Extended Data Fig. 1A**)^20^. NVEs tend to have low numbers of total junction reads per sample, which can make estimates of Ψ uncertain (**Extended Data Fig. 1B**). We therefore de-noised the estimate using a beta-binomial model. The workflow is described in **Extended Data Fig. 1C, Methods** and **Supplementary Note**. The method fits a smooth population Ψ distribution for the NVE across all individuals with data for a particular tissue, and generates credible intervals for all Ψ estimates.

After some filtering for NVE presence in at least one person and absence in at least one other person (see **Methods**), we identified 425,852 tissue/variably-spliced-region pairs across all 49 tissues in the GTEx dataset (**Supplementary Table 1**). Since many NVEs are observed in multiple tissues, this represents 56,415 unique NVEs, roughly balanced between alternative splice sites and skipped exon types (**Supplementary Table 2**).

Using these data, we define a new, interpretable summary statistic, exon frequency (EF), that describes the frequency of splicing of an NVE in a population, somewhat analogous to allele frequency (AF). The EF of a NVE in a given tissue is the proportion of individuals that splice the NVE at a threshold Ψ level or above. We use a 5% Ψ threshold because previous studies have suggested that this level of inclusion is near the lowest level where sequence conservation is commonly observed^19^.

Distributions of EF values using different Ψ cutoffs are shown in **Extended Data Fig. 1D-E**; 10% and 20% are generally similar while 1% likely introduces some noise. The steps in data processing to estimate EF for a given NVE are described in **Fig 1A**. EF is in some ways analogous to AF, but because estimation of EF depends on the availability of RNA-seq datasets from relevant tissues, the precise value is probably less important than the ability to stratify NVEs into rarer and more common subsets.

**Figure 1.**
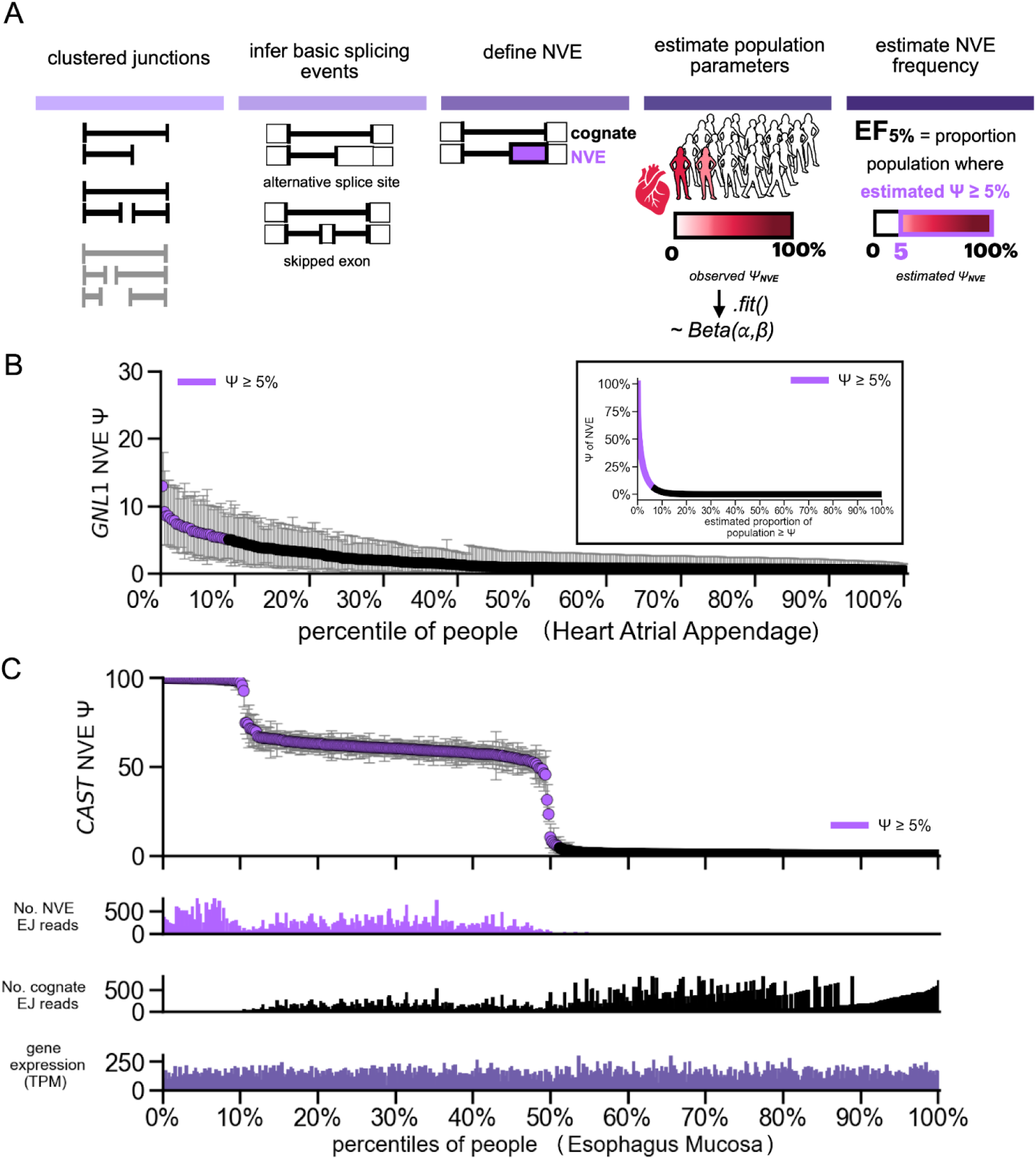
Overview of the method to detect NVEs and examples. A) Model figure describing NVE detection and our new summary statistic, exon frequency (EF). To define NVEs, we first filtered LeafCutter outputs for either alternative splice sites or skipped exons. For a given NVE observed in the RNA sequencing data, the percent spliced in (Ψ) of an individual can be directly estimated using the beta binomial model. The EF summary statistic is the percentage of individuals whose Ψ exceeds a given threshold, shown for a 5% Ψ. B) An example exon in *GNL1* with EF_5% Ψ_ = 0.1 in a heart tissue. Individuals with heart tissue samples are sorted by their estimated Ψ values for this exon, which is shown as percentiles on the bottom x-axis. The mean posterior estimates on Ψ is plotted (highlighted in purple if the estimate is at least 5% Ψ in the individual), along with error bars showing 95% credible intervals on Ψ. In the inset, an example of the population distribution used to calculate the exon frequency (EF) of this exon. Note that the y-axis scales differ between plots. C) An example of an NVE in *CAST* gene (with EF_5% Ψ_ = 0.6) that has high variability amongst individuals. Individuals with Esophagus Mucosa data are sorted by their estimated Ψ values for this exon, which is shown as percentiles on the x-axis. Top plot: The mean posterior estimates on Ψ are plotted (in purple if the estimate is at least 5% Ψ in the individual), along with error bars showing 95% credible intervals on Ψ. Below: bar plots of EJ reads for both NVE and cognate, and overall observed mRNA levels (TPM). Note that the y-axis scales differ between plots.

To illustrate our approach, we provide an example of a fairly typical NVE, which occurs in the *GNL1* nucleolar GTPase gene and has an EF of 0.1 (**Fig. 1B**). This exon appears completely absent or very lowly included in most people, but occurs at a Ψ of 5-10% in about 10% of individuals, with credible intervals on Ψ well separated from zero. This splicing variation does occur against a background of fairly uniform gene expression (**Extended Data Fig. 2A**). The underlying factors that drive splicing differences between individuals might include genetic variation acting in *cis* or in *trans*, or environmental factors, and are not clear in this particular case. Examples of NVEs at different EFs are shown in **Extended Data Fig. 2B-D**. Examining the splice site motifs of NVEs, we observed that they match consensus motifs to a similar degree as the splice sites of cognate exons (**Extended Data Fig. 2E-F**), similar to what has been seen for alternative exons overall ^21^. We provide various summary statistics describing the splicing of NVEs, such as median and maximum Ψ across the spectrum, in **Extended Data Fig. 3 and 4**, observing positive relationships between most measures of exon inclusion and EF (**Extended Data Fig. 4C-D**). NVEs can even be constitutively spliced in some people, and absent from others, as shown by a second example, occurring in the calpastatin gene *CAST* (**Fig. 1C**). These examples highlight diversity in the splicing patterns of exons across individuals.

We next sought to understand the distribution of NVEs across genes and individuals. Since many NVEs are detected in multiple tissues, we focused on the tissue where the exon achieves its highest EF, which we call “top EF” and use as the default EF value for an NVE. Using our approach, we identified 28,675 alternative splice site NVEs (NVE_alt ss_) and 27,740 skipped exon NVEs (NVE_se_) in all of GTEx. We find that NVEs are widespread, occurring in 74% of protein-coding genes, with a median of 3 NVEs per gene (**Fig. 2A**). Across all 49 tissues, some variability was observed in the number of NVEs, with a median of ∼2200 NVEs per tissue (**Extended Data Fig. 5A**).

**Figure 2.**
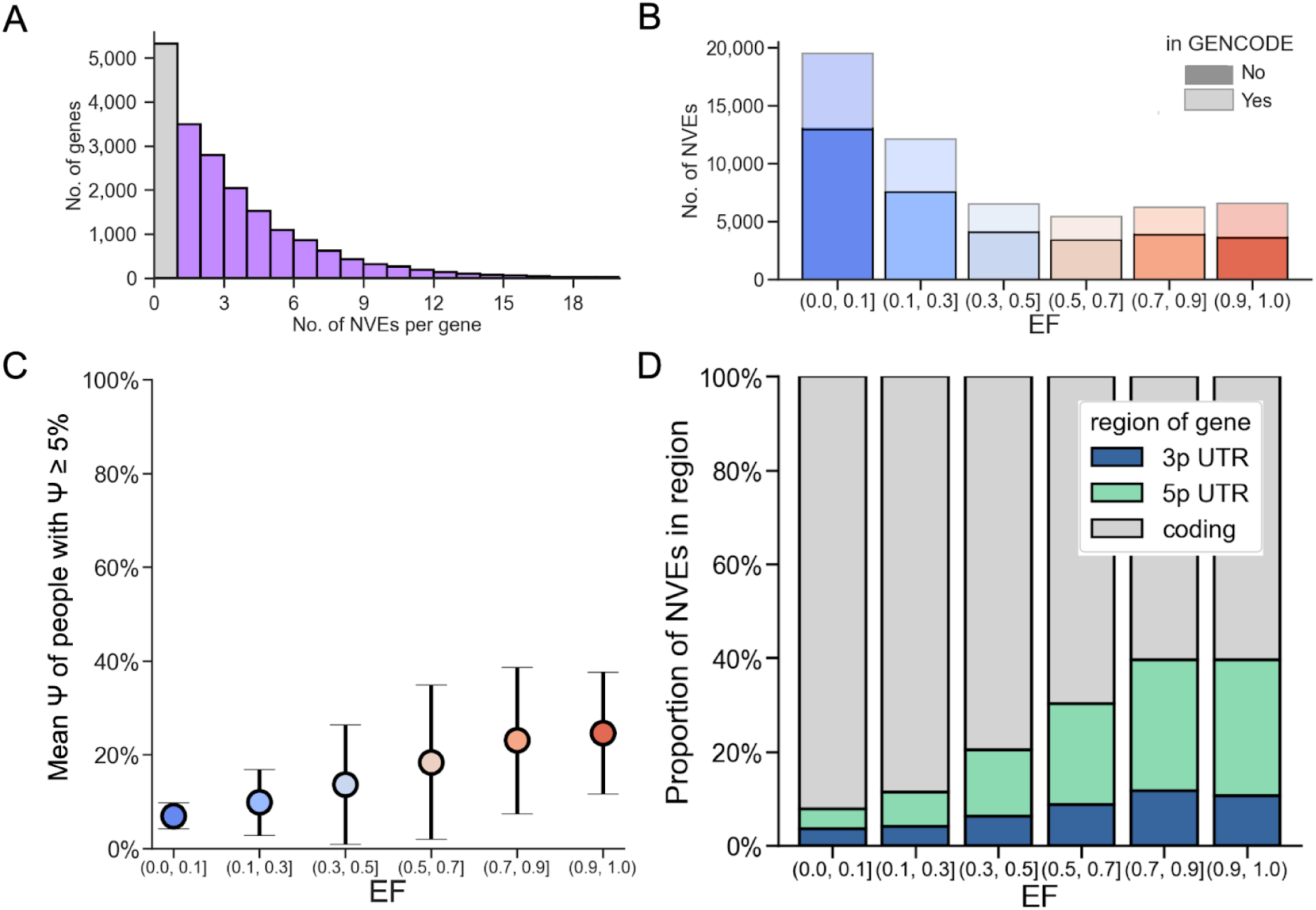
The distribution of NVEs across genes, gene regions, and their relative usage in individuals. A) Histogram of the number of NVEs in a given gene, including those protein coding genes with no NVEs (gray) at least one NVE (purple). B) The top EF spectrum in GTEx. For each NVE, the tissue with the most usage of the NVE was taken, indicating the highest possible EF in GTEx. We use this color scheme throughout the paper with lower EFs shown in blue and higher EFs shown in red. Lighter boxes indicate that the NVEs are not on GENCODE reference annotations. C) Average ψ of NVE in individuals who splice in ≥ 5% ψ. 1000 randomly sampled NVEs were selected, repeated 10 bootstraps. Using the alpha and beta parameters of the population distribution for each NVE, we estimated the integral for the region of the beta distribution where ψ was predicted to be ≥ 5%. We used scipy integrate.quad to estimate the integral. Mean estimates shown with mean of standard deviations across the 10 bootstraps. D) Proportion of NVEs in given gene regions, split by exon frequency.

We estimated the average number of NVEs expressed per individual as ∼9,000, using a conservative approach summing the EFs from a top EF set calculated from the ten best-sampled tissues. Although low-EF NVEs are more numerous, those with higher EFs are more commonly observed, so the EF distribution of the set of NVEs that occur in any given individual is will be skewed toward higher EF values (**Extended Data Fig. 5B**). As a result, the fraction of NVEs shared by any two (unrelated) individuals – calculated as the sum of the squares of EF values over all NVEs over the sum of the EFs – is just over half (∼58%).

### Most NVEs occur in coding regions and have low EF

We found that the distribution of EF values of NVEs is skewed toward lower values, with most NVEs being rare in the population, but some being very frequent (**Fig. 2B**). We estimate that the mean ψ of an NVE in individuals that splice above our threshold level (5%) ranges from ∼10% for low-EF NVEs (top EF 0.1 or below), to just over 20% for high-EF NVEs (top EF 0.9 or above) (**Fig. 2C**). The RNA-seq read depths available in GTEx are more than adequate to distinguish these moderate ψ values from absence of splicing, and NVEs possess splice site motifs very similar to those of other exons (**Extended Data Fig. 2D**), even for low-EF NVEs (**Extended Data Fig. 2E**).

We explored the features associated with EF for NVEs. As noted above, more highly included exons (higher median ψ) tended to have higher EF values (**Extended Data Fig. 4C-D**). For NVE_alt ss_, their 5’ and 3’ splice site (SS) motif scores (using MaxEnt^22^) tend to be slightly weaker than their associated cognate SS, with the difference shrinking at higher EF values (**Extended Data Fig. 5C**). These observations suggest that the SS strength of NVEs is a contributor to their inclusion and EF values.

Notably, 60% of NVEs present in GTEx are absent from comprehensive reference annotations (GENCODE 45). As expected, NVEs with low EF are predominantly unannotated, while around half of high-EF NVEs are annotated **(Fig. 2B)**. Low-EF NVEs that are absent from reference annotations have more tissue-restricted splicing than those that are present in reference annotations (**Extended Data Fig. 5D)**. Different NVEs were restricted to different subsets of tissues, with no prominent outlying tissues (**Extended Data Fig. 5E**). In total, NVEs expand the sequence space covered by the transcriptome by ∼1.4 MB (**Extended Data Fig. 5F**). The number of NVEs detected in an individual increases with read depth (**Extended Data Fig. 5G**), suggesting that substantially more NVEs could be found with deeper sequencing.

In general, alternative exons occur most commonly in coding regions, but also occur fairly often in 5’ UTRs and rarely in 3’ UTRs of genes. We found that most NVEs occur within the region of the gene that contains coding exons, a class we call cdsNVEs, and which tend to have low EF values. A substantial minority of NVEs occur in 5’ UTRs, and this subset tended to have higher EFs (**Fig. 2D**). This observation is consistent with previous work showing that evolutionarily new alternative exons arise most commonly in 5’ UTRs^8^. NVEs rarely occur in 3’ UTRs, likely because they contain very few introns relative to other transcript regions. In summary, both splice site strength and location in the gene appear to contribute to the emergence and/or maintenance of NVEs in the human population.

### NVEs impact mutationally-constrained genes

To explore the evolutionary properties of genes with and without NVEs, we considered gene tolerance to germline LoF mutations in the human population, which is a proxy for both gene function and selection^23^. This property has been quantified across nearly all human genes by computing the loss-of-function observed to expected upper bound fraction (LOEUF) across a population, where lower scores correspond to more constraint (greater intolerance to LoF) in the gene^23^.

Surprisingly, we find that the distribution of LOEUF scores are lower for NVE-containing genes than genes which lack NVEs (**Fig. 3A**), even after adjusting for the number of annotated exons per gene (**Extended Data Fig. 6A**). Furthermore, the distribution of EFs was shifted toward lower values in more constrained genes (**Fig. 3B**). Such a pattern could occur if NVEs arise more commonly in more constrained genes but often experience selection favoring lower ψ values, driving EFs lower, although other scenarios are possible. NVEs in less-constrained genes tend to have higher EFs, some of which might become canonical alternative exons over evolutionary time. Genes containing canonical alternative exons – which are spliced in all or virtually all people – showed no bias in LOEUF score relative to all genes (**Extended Data Fig. 6B**).

**Figure 3.**
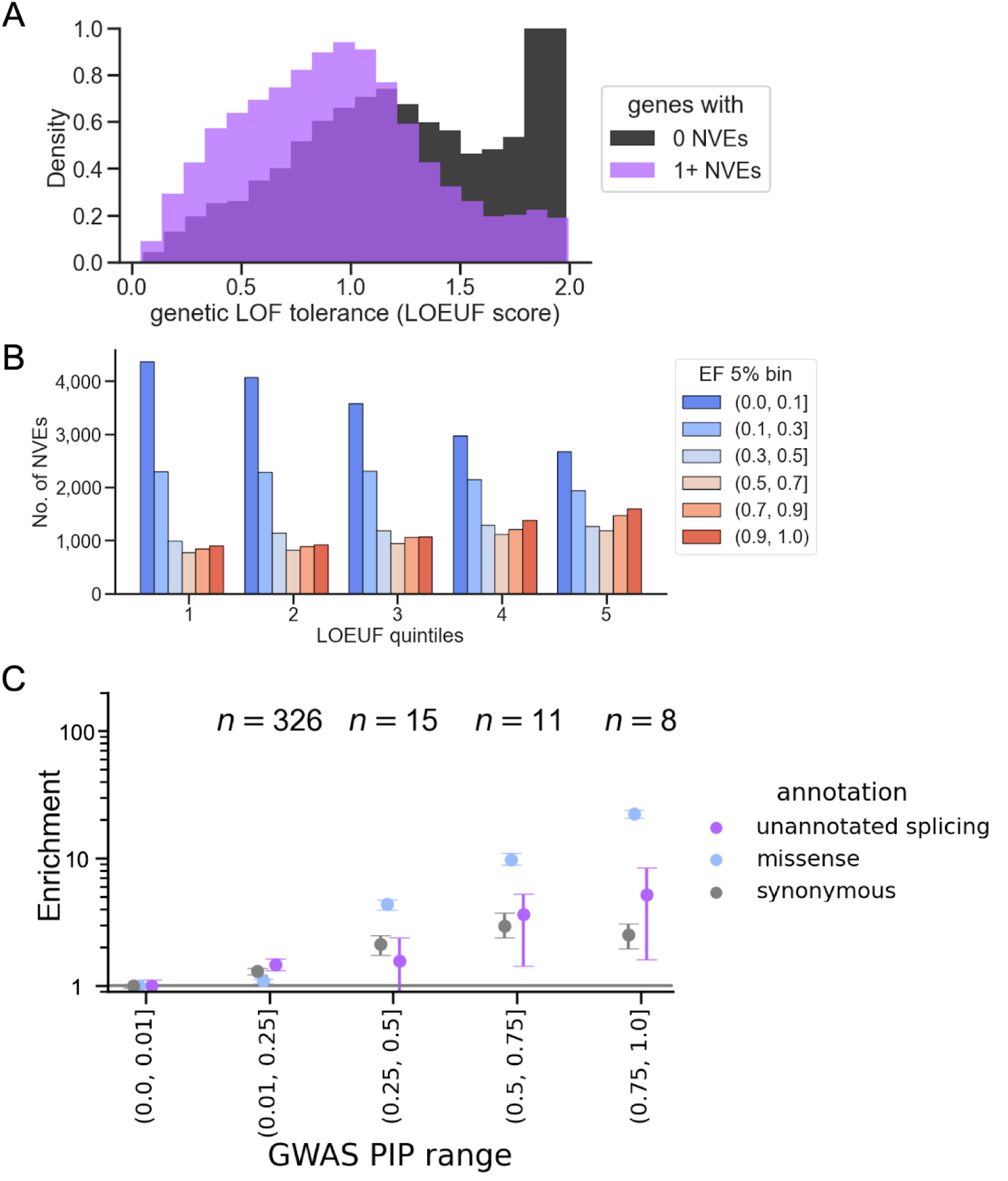
NVEs tend to impact more mutationally constrained genes, and provide additional functional interpretations in GWAS. A) Distribution of LOEUF scores of genes with NVEs (purple) and genes that do not have any NVEs (black) Significant by KS test p-value 10^-217^. We dropped duplicates of genes that had multiple NVEs. B) The percentage of genes stratified by NVE EF bin, represented in each LOEUF quintile, with low values indicating more constraint. Genes that contain low EF NVEs are enriched among the most constrained genes, whereas genes with high EF NVEs are largely unconstrained. Here, the EF represents the NVE that is most frequent across the gene. C) GWAS from three global biobanks (UKBB, FG release 9, and BBJ) were pooled together. We included NVEs that were present in all individuals to boost power. Unannotated splicing events are enriched above synonymous at high PIPs. For the unannotated splice site set, pLOF, annotated splice region, nonsense, and missense variants were filtered out of this set to ensure enrichment was not driven by well-explained variants. Number of variants being categorized as within an unannotated splice site is shown as n values.

### NVEs aid interpretation of GWAS variants

To explore whether NVEs could improve disease interpretation, we focused on variants that occur in the extended splice site motifs of NVEs not present in GENCODE comprehensive reference annotations. We analyzed the results of statistical fine-mapping of GWAS for around 1300 traits across three global biobanks: UK BioBank, FinnGen, and Biobank Japan (UKBB, FG, BBJ, respectively)^24^. Statistical fine-mapping yields a posterior inclusion probability (PIP) for each variant, which reflects the probability that the variant causally drives the association at the locus and enables enrichment analyses of fine-grained annotations, such as NVEs, that are not powered for heritability-based analyses (**Supplementary Note**).

The observed enrichments of causal variants in NVEs are above the level of synonymous variants, particularly at high PIP values, but below those of missense annotations (**Fig. 3C**). To ensure that the enrichments were not driven by existing annotations, the enrichments shown exclude GWAS variants already annotated as genetic LoF, splice region, or missense variants. We included NVEs that were present in all individuals to boost power. Performing enrichment analyses separately for each biobank reduced power, but still had a similar trend of effect (**Extended Data Fig. 6C**).

While synonymous variants are typically null or nearly null in analyses of ultra-rare variants^25^, some synonymous variants have been identified as high-confidence causal variants in cross-biobank fine-mapping^24^, and may cause changes in splicing, mRNA stability or translation, for example, so the enrichment for synonymous variants in high-PIP bins is unsurprising.

We next explored the utility of unannotated NVEs for GWAS interpretation. One example is a pleiotropic synonymous variant in *ASGR1* (rs55714927, MAF 15%) that is both a sQTL and an expression quantitative trait locus (eQTL) and colocalizes with an NVE (EF 7%) splice site. This common *ASGR1* variant is known to increase the inclusion of this unannotated NVE, and this exon contributes to phenotypes such as cholesterol and heart function^24,26,27^. We found that, in many cases, the alternate allele was associated with increased splicing of the NVE to between 3 and 10%, while individuals with the reference allele had NVE Ψ values near 0% (**Extended Data Fig. 6D**). We observed a negative correlation between the Ψ of this NVE and the expression of *ASGR1* (**Extended Data Fig. 6D**), consistent with the potential of this exon to trigger NMD. This example illustrates a molecular mechanism for a synonymous variant and demonstrates that even NVEs with low Ψ and EF values can be relevant to GWAS loci.

### Low observed Ψ NVEs can have outsized impacts on expression

To explore the effects of NVEs on coding potential and expression more broadly, we built a custom pipeline to identify cdsNVEs that have NMD potential, including causing a frameshift or introducing a stop codon (Methods), which we refer to as nmdNVEs. nmdNVEs had low Ψ values, which were lower on average than cdsNVEs without NMD potential (**Extended Data Fig. 7A**).

We considered whether splicing of nmdNVEs tends to reduce gene expression. In order for an NVE to meaningfully influence gene expression, it must be spliced in to a sufficient extent to reduce the level of the cognate isoform. Degradation of the NVE-containing isoform by NMD is expected to reduce both the Ψ value that is observed in RNA-seq data (which reflects levels of mature mRNAs), and also reduce the final expression level (**Fig. 4A**). Because of the impact of NMD, even a relatively modest observed difference in cytoplasmic PSI value can be predictive of a fairly large change in gene expression. In the hypothetical example illustrated, destabilization of the NMD isoform by 5-fold relative to the canonical isoform^28^ implies a nuclear PSI of 62% and a 2-fold reduction in gene expression associated with an exon whose observed PSI is just 25%. Under the same assumptions, a frame-disrupting exon with an observed PSI of 10% would reduce expression by ∼29%.

**Figure 4.**
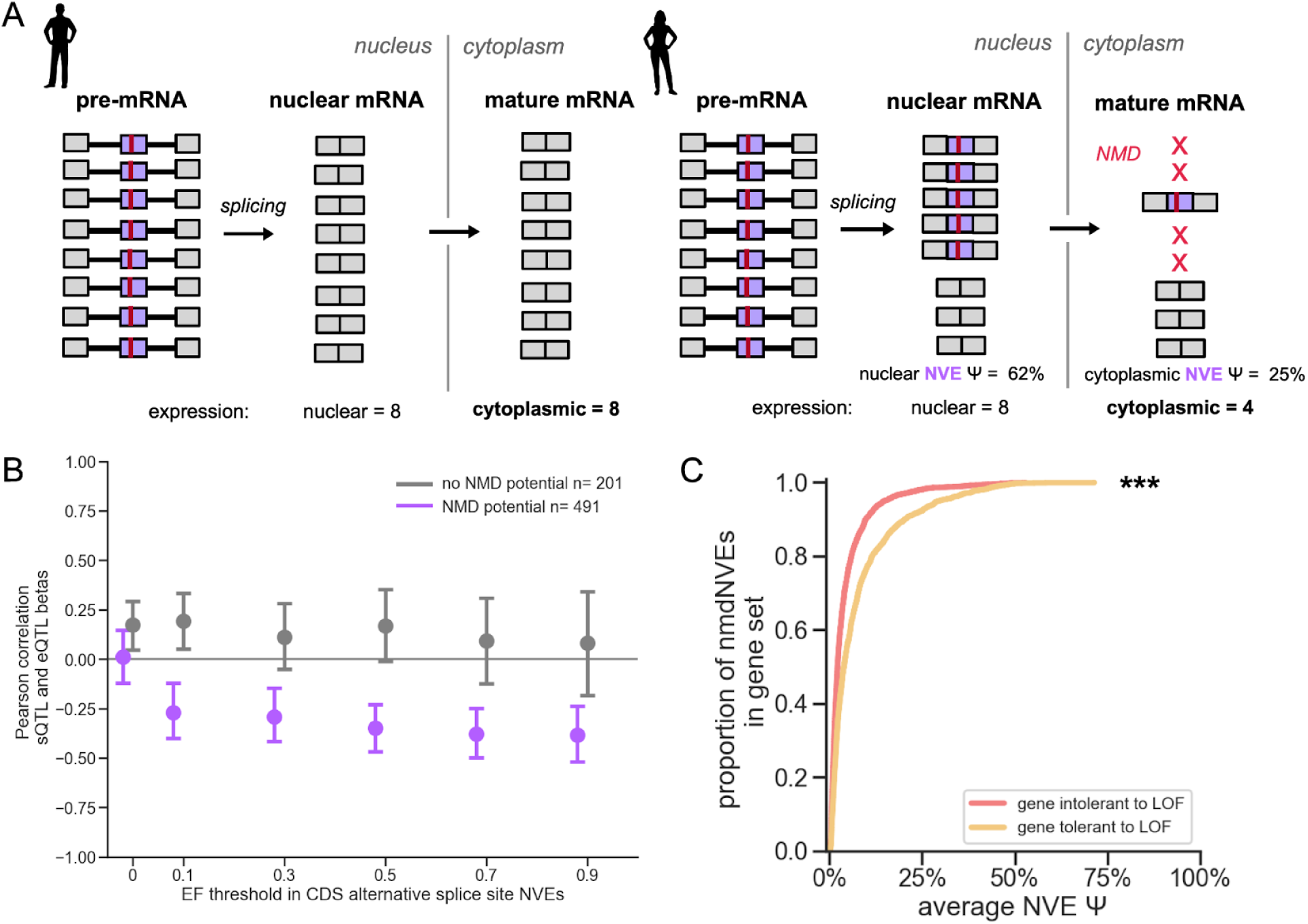
Evidence of gene expression changes of NVEs with NMD-potential across the EF spectrum. A) Lowly spliced NVEs can have outsized impacts on expression. Left individual: the NMD-causing NVE (purple, with red bar for stop codon) is not spliced into transcripts, and will have no impact on expression. Right individual: Inclusion of the nmdNVE in ⅝^ths^ of transcripts in the nucleus is implied by the Ψ of 25% in the cytoplasm, assuming that the stability of nmdNVE isoforms is reduced 5-fold by NMD, resulting in a 2-fold reduction in gene expression. In general, if NMD reduces stability of nmdNVE isoforms by *k*-fold of an nmdNVE with cytoplasmic Ψ value of *x*, gene expression will be reduced by (*k*–1)*x*/(1+(*k*–1)*x*). B) Effect size estimates (slope of regression of QTL) shown for genetic variants that were nominally significant sQTLs for NVEs, that are also significant eQTLs in the same gene in the same tissue. Genetic variants were filtered out if in complex loci (i.e. were eQTLs in other genes in the tissue). Note that association between variant and splicing event is with the cognate splicing event, such that a decrease in splicing would imply an increase in the overall NVE:cognate ratio, so we multiplied the effect size by –1. cdsNVEs are further separated into those with NMD potential (purple) and without (black). Shown are mean and range (2% to 98%) of 1000 bootstrapped Pearson correlations between sQTL and eQTL effect sizes, performed at increasing EF thresholds. C) The average Ψ across individuals for a given NVE was estimated in the tissue. Showing in this plot any EF NVE that causes NMD, further separated depending on if the NVE was unconstrained (top 10% percentile of LOEUF) or constrained (bottom 10% of percentile LOEUF) gene. *** Indicates significant KS test (10^-43^) between the two groups.

Sufficient read depth is therefore required to both detect the observed splicing of the NVE, and to observe any associated change in expression. Because of these considerations, we chose to explore this question using a subset of NVEs that have large variability in splicing, occur in genes with variable expression, and have sufficient read depth for both types of variation to be observed. Specifically, we considered NVE_alt_ss_ whose splicing is significantly associated with a genetic variant that impacts both splicing and gene expression, i.e. that is both an sQTL and an eQTL in GTEx. In this set, we observed a negative relationship between inferred NVE sQTL effect size and (directional) eQTL effect size for nmdNVEs, and this relationship is consistent across nearly all EF thresholds (**Fig. 4B**). This observation supports the idea that, even at low observed Ψ values, nmdNVEs often reduce expression by shifting the mRNA output of a gene from productive, stable mRNAs to unproductive/unstable mRNAs. By contrast, no significant relationship between sQTL and eQTL effect size/direction was observed for cdsNVEs that are not nmdNVEs. Such NVEs presumably yield unique protein isoforms, though this direction was not explored here.

Because many NVEs occur in annotated introns, they may create non-functional mRNA and protein isoforms (whether they trigger NMD or not), which might often be mildly deleterious, particularly in constrained genes. We observed that in constrained genes, nmdNVEs tend to have lower Ψ values than nmdNVEs in unconstrained genes (**Fig. 4C**), likely reflecting selection against large perturbations of gene expression. Notably, for NVEs in 5’UTRs, higher Ψ values were associated with increased gene expression (**Extended Data Fig. 7B**). NVEs in this gene region may positively impact gene expression via exon-mediated activation of transcription starts (EMATS)^29^.

Together, the observations above indicate that the splicing of NVEs can be driven by genetic variants such as SNPs, that NVEs may impact gene expression negatively or positively, and that, in general, selection may favor reading frame-preserving NVEs and NVEs that exert smaller effects on gene expression over other NVEs.

### Both genetics and age impact the presence of NVEs

Splicing can be impacted by both genetic and environmental factors^11,30^. To understand the basis of inter-individual differences underlying differences in NVE splicing, we asked whether common genetic variants or other factors were associated with presence of NVEs. We find that 60% of NVEs are associated with at least 1 sQTL (defined as a variant within a credible set in the fine-mapping dataset) in their top EF tissue, suggesting that many NVEs are modulated by *cis*-acting genetic variants. This proportion varied only modestly across EF bins (**Fig. 5A**). This pattern is consistent with prior work showing that splicing variation in GTEx is mostly driven by *cis* genetic effects^31^.

**Figure 5.**
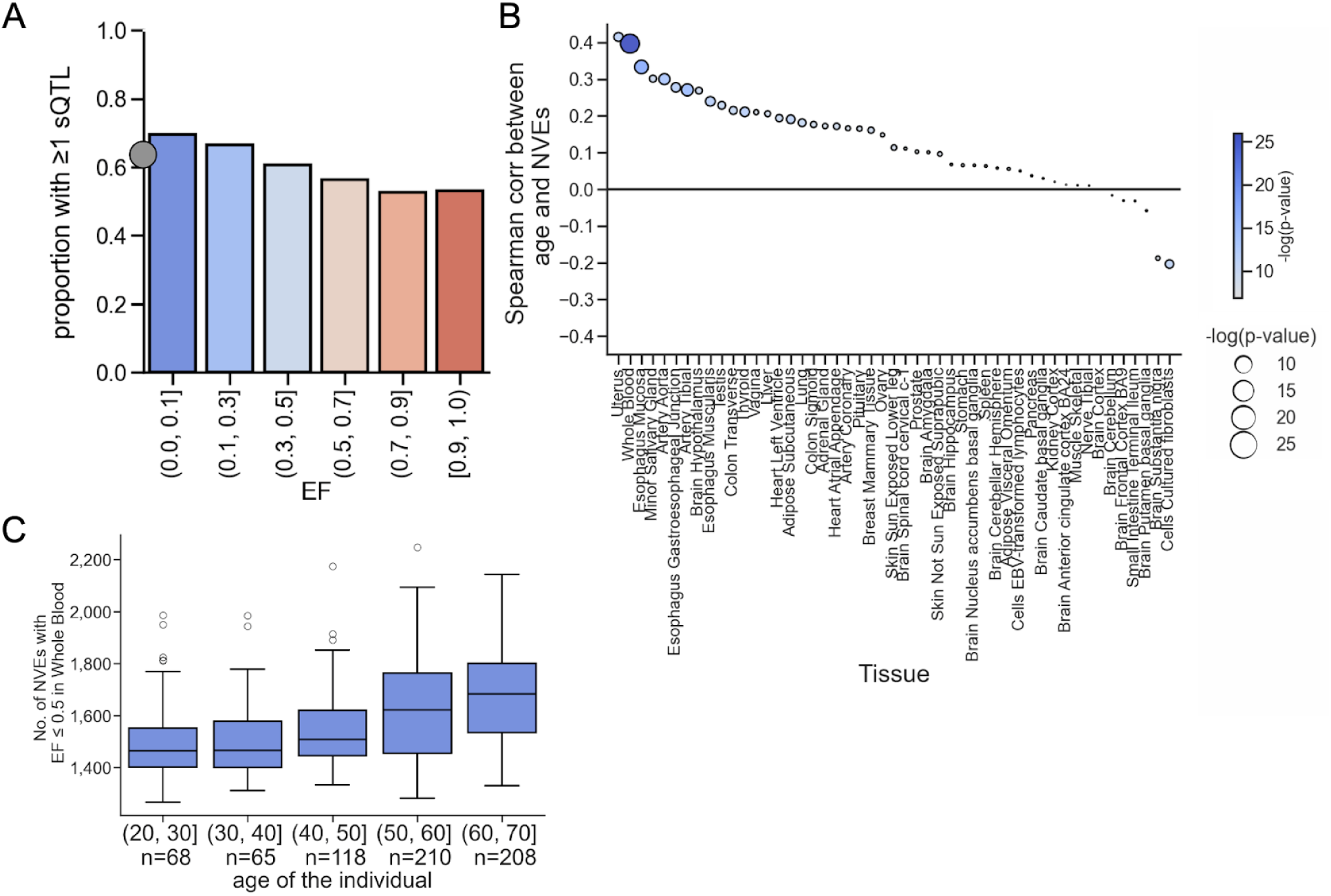
Both genetic and non-genetic factors contribute to splicing of NVEs. A) Fraction of NVEs in each EF bin explained via ≥1 *cis* sQTL in GTEx. Gray dot indicates the average across all NVEs. B) A Spearman correlation between age of individuals versus moderately mid-to-low frequency NVEs (≤0.5 EF) across all tissues was performed across all tissues. Here we show the correlation across all tissues, with both the color and size of dot indicating the -log_10_ p-value of the correlation. Strip plots of Bonferroni-corrected marginally significant tissues are provided in **Extended Data**. C) Boxplot of number of NVEs versus age of individual in most significant p-value tissue from B; Whole Blood.

Beyond genetic factors, we asked whether the age of the individual impacted NVE splicing, as age has been associated with splicing changes^32–35^. We found that the number of NVEs was positively correlated with age of the individual in many tissues (**Fig. 5B, C**). These correlations appeared to be driven mostly by increased NVE usage over the age of fifty. These observations suggest that splicing fidelity might decrease with age in many tissues, or that selective pressures on gene expression might loosen with age. Data for selected tissues are shown in (**Extended Data Fig. 8A**). The primary exception to this pattern was for cultured fibroblasts, which yielded a significant negative Spearman correlation (10^-6^) between age and NVE count. This difference in age-dependence might relate to changes in age-associated epigenetic state induced by cell culture^36^. We also show that age is not correlated with read depth (**Extended Data Fig. 8B**).

### Splice modifier score uses NVEs to improve common variant interpretation

The identification of variants that impact splicing remains challenging. sQTLs directly correlate common genetic variants with splicing measurements, but because of pervasive linkage disequilibrium they do not directly identify which variant causally impacts splicing, and do not inform about mechanism. Fine-mapped sQTLs are a gold standard, but because of limited statistical power, sets of fine-mapped sQTLs are far from complete. For rare variants (MAF < 1%), sQTLs and fine-mapped sQTLs are even less powered. These considerations motivate the importance of predicting splice-affecting variants using other sources of information.

SpliceAI, a deep neural model trained on human GENCODE annotations, is a widely-used tool for prediction of splice-impacting variants based on primary sequence^16^. However, this method, and methods built on top of it, such as ABsplice^17^, have been less successful in predicting effects of common variants, which often have low effect sizes relative to rare variants. We therefore attempted to improve identification of more common splice-affecting variants.

For the prediction task of finding common splice-affecting variants, we also considered two distinct situations: the case in which in-sample RNA-seq data is available, and the case where only reference data are available. To this end, we developed the Splice Modifier Score (SMS), analogous to the Expression Modifier Score (EMS) for eQTLs^37^, which uses a regression model to estimate the probability that a variant modifies splicing. We trained SMS using fine-mapped sQTL data^38^, using highest probability sQTLs (PIP of 90% or above) as positives, and low-probability sQTLs (0.2% or below to improve class balance) as negatives. We consider the top association of a variant that is within 5 kb of a given splicing phenotype in a tissue, considering skipped exons and alternative splice sites in protein-coding genes.

To train the model, we annotated genetic variants with a set of molecular features including: distance to nearest splice site, known exonic splicing enhancer^39^ and silencer^40^ motifs, splicing-associated histone marks^41^, and binding sites of RNA-binding proteins based on eCLIP data^42^. We trained SMS on 80% of sQTLs (randomly selected) and used the remaining data as a held-out test set, yielding stable performance across many trials. When given access only to reference gene annotations, where the distance feature uses GENCODE splice sites (SMS-GENCODE), the method’s performance improves moderately over SpliceAI (**Fig. 6A**). However, when using a distance feature based on GTEx splice sites, including those belonging to NVEs, our “SMS-full” model achieves an area under the precision recall curve (AUPRC) of 0.49, substantially better than SpliceAI or SMS-GENCODE **(Fig. 6A**). When training on the top splicing signal (top PIP), i.e. the best splicing association for each unique variant, we obtained qualitatively similar results and the same ranking of algorithms (data not shown).

**Figure 6.**
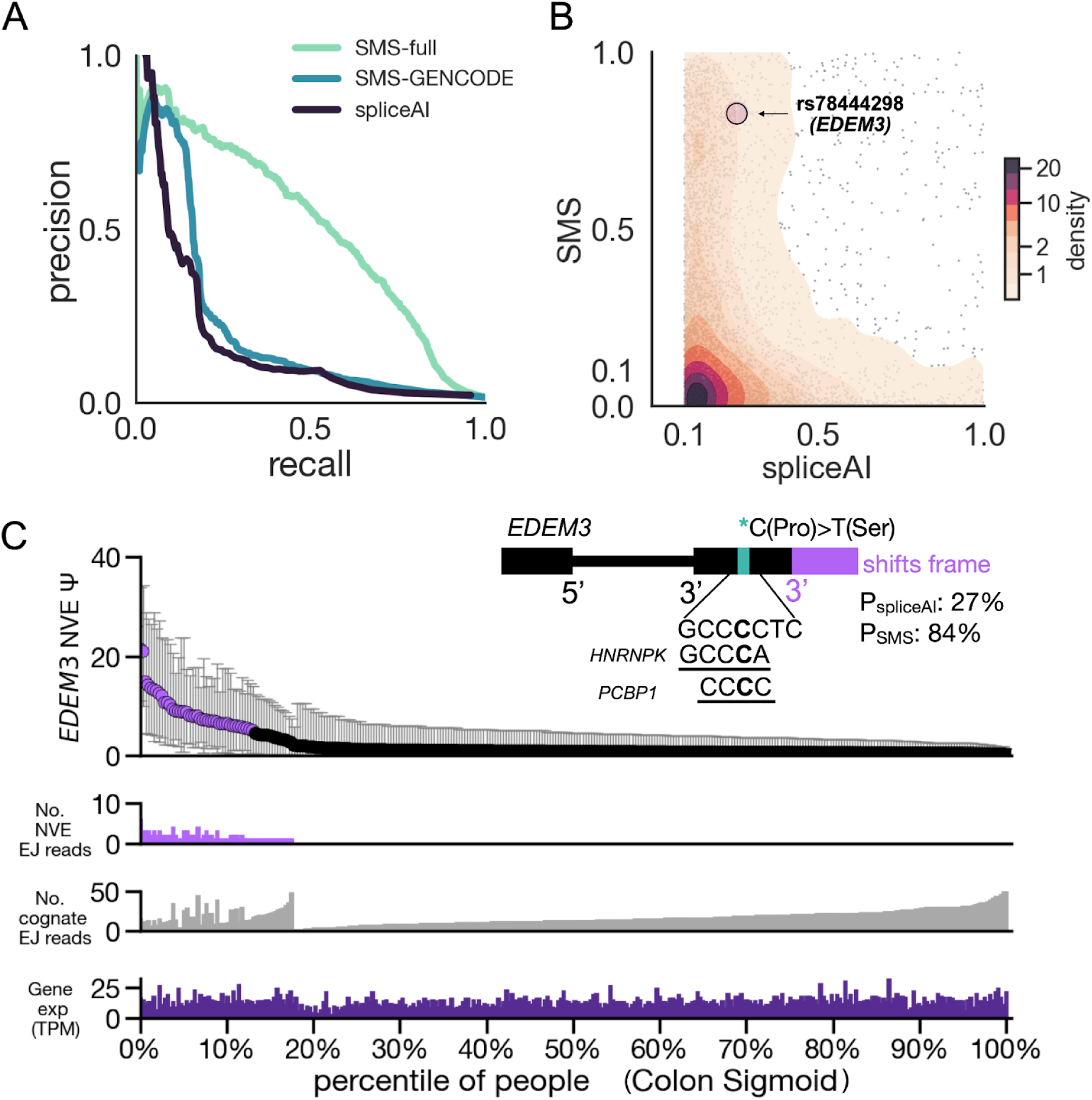
Splice modifier score (SMS) provides insight into variant induced splicing and helps interpret GWAS. A) Comparison of model performance by precision recall curves (PRC) on a held out test set. PRC of full logistic regression model trained on GTEx distances, PRC of full logistic regression trained on GENCODE distances only, and PRC of logistic regression trained only on SpliceAI score is shown. B) Comparison of SpliceAI and SMS scoring. SMS was scored using full model parameters and distance to GTEx splice sites across all GTEx variants. Variants with SpliceAI that scored >0 were selected, and relative SMS and SpliceAI scores were compared in a scatter and kernel density plot. High density of points along edges suggest that SpliceAI and SMS are picking up orthogonal information. Colors indicate relative density. C) Missense variant in *EDEM3* protein (rs78444298) is associated with many GWAS studies and may cause a NMD-inducing NVE in GTEx. Variant is a negative sQTL in GTEx for the cognate exon of the NVE (meaning a positive sQTL), and negative eQTL in GTEx for the same gene (see GTEx browser). Individuals with artery tissue samples are sorted by their estimated Ψ values for this NVE, which is shown as percentiles on the bottom x-axis. The mean posterior estimates on Ψ is plotted (highlighted in purple if the estimate is at least 5% Ψ in the individual), along with error bars showing 95% credible intervals on Ψ. Dot is colored purple is estimated Ψ exceeds 5%. Also showing bar plots of exon junction (EJ) reads for both NVE and cognate, and overall observed mRNA levels (TPM). Inset: Details on this variant. rs78444298 induces a proline to serine mutation (shown in diagram) and occurs near a 3’ splice site. The NMD-inducing mechanism is also shown: low EF NVE boundaries from an alternative 3’ SS with a maxent score of 10 (purple). Two out of the four RNA binding sites predicted to bind to the region (from oRNAment database) are highlighted, showing the two RNA binding proteins where the motif clearly is losing binding due to the variant presence. Probabilities of variant causing splicing changes using SpliceAI and SMS predictors shown in teal blue.

Examining the performance of models using different combinations of features, we observed that distance to nearest GENCODE splice site contributes to precision, analogous to studies of eQTLs where distance to the reference transcription start site (TSS) is highly predictive^43^, but that use of additional information related to splicing is needed to achieve a high AUPRC **(Extended Data Fig. 9A)**. Together, these observations support that the improved accuracy of SMS-full over the SMS-GENCODE model is driven by inclusion of unannotated splice junctions, such as those of NVEs, and demonstrates the importance of in-sample RNA sequencing to better predict the genetic basis of splicing variation. Overall, we found that ∼30% of splicing phenotypes associated with sQTLs are NVEs.

Comparing SMS-full to SpliceAI predictions for individual variants, we observed substantially different scores for the two methods in many cases (**Fig. 6B**), suggesting the potential complementarity of these models (**Extended Data Fig. 9B**). One explanation for the large divergence between predictions could be that SpliceAI performs best on rare variants that induce large changes in splicing^16^, while SMS-full is trained on data including more subtly spliced NVEs, potentially improving its ability to identify common variants which tend to exert smaller effects.

An example of a variant with low SpliceAI probability (0.27) but high probability with SMS (0.94) is rs78444298, located in EDEM3 with MAF 1.5%. This variant is associated with schizophrenia, as well as metabolic and blood phenotypes^44–46^. Molecular studies of this variant have focused on EDEM3 as a LoF phenotype^47^. Though this is a predicted missense variant causing proline to serine change, whether LOF occurs at the level of RNA abundance or protein function remains unclear. In several GTEx tissues, the variant is associated with an increase in splicing of a low EF nmdNVE, also associated with decreased gene expression in several of those tissues. SMS indicates that the variant is in a region of potential binding by several RBPs (HNRNPK, PCBP1, SRSF5, TAF15) near the 3’ splice site of the nmdNVE. The variant alters predicted affinity for two of these RBPs (HNRNPK, PCBP1) by 5-fold and 4-fold, respectively, highlighting a potential molecular mechanism for this variant. The cognate exon and nmdNVE have moderate to high 3’ splice site scores (6.7 and 10.4 bits, respectively). In summary, the SMS model and NVE information provides novel molecular explanations for this variant. SMS scores for all GTEx variants are provided (**Supplementary Table 3**).

## Discussion

To better understand the impact of inter-individual variation in gene and isoform expression, we defined and identified over 50,000 NVEs, and characterized their impacts. Many variants identified in GWAS studies lie within noncoding regions, some of which impact pre-mRNA splicing. Here we found that NVEs tend to occur in more constrained genes, and show that NVEs enhance our ability to interpret GWAS variants, especially in the UK BioBank. As RNA-seq data become more available from GWAS participants, detecting NVEs from these datasets may further enhance interpretation for GWAS variants.

NVEs have potential to affect gene function or expression, commonly occurring in coding exons or their intervening introns, 5’UTRs, and less commonly in 3’UTRs. NVEs that occurred at higher frequency in the population were particularly enriched in 5’UTRs, parallelling the high frequency of evolutionarily new exons observed in this gene region^8^. NVEs with lower EFs occurred largely in coding regions, particularly in constrained genes, and typically had lower Ψ values. Natural selection may more strongly limit which NVEs can rise to higher PSI and EF values in coding regions, because of constraints on the expression levels of canonical isoforms. One limitation of our current study was we were only to investigate a small number (491) of eQTL- and sQTL-associated NVEs with NMD potential (**Fig. 4**). These NVEs, whose splicing impacts gene expression by producing nonfunctional isoforms, could be targeted with splice-switching ASOs to therapeutically increase (or decrease) protein abundance^48,49^. Inhibiting the splicing of an NVE would only be useful in people that splice the NVE, of course, but activating the splicing of an NVE via an ASO might be feasible even in individuals where the NVE is not normally included.

Recent efforts have begun to more comprehensively describe the impacts of inter-individual variation on splicing^11,31,50^. In principle, NVEs might arise from cis-acting mutations that create or disrupt splice sites or splicing regulatory elements, from trans-acting mutations that impact the activity of splicing factors, or from various physiological or lifestyle factors such as sex, age, diet, environmental exposures, etc. Here, we find strong evidence that a majority of NVEs are impacted by cis-genetic variation (Fig. 5A). Trans-acting variation is also likely to be important but can be more difficult to detect. Sex-specific differences in splicing are also known^51,52^, and there are widespread splicing differences in individuals with certain diseases, including autism spectrum disorders^3,53^. Some of these differences have been shown to be modulated by specific splicing factors^3,53^. Age also influenced NVE occurrence, although this effect varied by tissue (**Fig. 5B**).

All humans are thought to express very similar sets of genes, in similar tissue-specific patterns^7^. However, we found here that individuals commonly differ in the use of specific exons, with each person expressing several hundred NVEs in each well-sampled tissue. Comparing two random, unrelated individuals, just over half of NVEs are expected to be shared, implying that unrelated people differ in the presence of hundreds of different mRNA isoforms in each of their tissues. Our findings using SMS emphasize the importance of using diverse and population-relevant RNA-seq data for the inference of genetic variation that modulates splicing. The preponderance of lower-EF exons and their enrichment in constrained genes, which are more associated with disease, emphasizes the value of increasing population sample sizes rather than simply increasing read depth for increased detection of disease-relevant splicing.

## Author contributions

HNJ performed the analyses and authored the manuscript. BLG helped implement the partial pooling analyses, and provided feedback on the manuscript. JG helped with construction of NMD analyses with guidance from HNJ. MK provided guidance on GWAS results. HKF, KJK, and CB provided guidance on analyses, feedback on manuscript, and helped guide project directions.

## Acknowledgements

We thank Mike McGurk for guidance on analyses, feedback on the early manuscript, as well as beta-binomial model framework. We thank Jacob Ulirsch, who along with Masahiro Kanai, introduced the *ASGR1* exon. We also thank Ran Cui for providing guidance on fine-mapping based analyses. We thank Chi-Hsien Chang for help testing the code. We thank Athma Pai, Kaitlin Samocha, and David McWatters for direct feedback on the manuscript. This work is supported by the Novo Nordisk Foundation. (NNF21SA0072102).

## Methods

### Ethics statement

No primary data were generated for this study. Person-related data were obtained through authorized access from primary data controllers.

### Statistics and reproducibility

No statistical method was used to predetermine sample size. We did not use any study design that requires randomization or blinding. In the GTEx data no samples were excluded.

### Datasets

#### GTEx release v8

From the GTEx download data portal, we downloaded splicing “phenotype” files, which were originally used as inputs of the original sQTL study in GTEx (v8p hg38). We accessed it using the <u>GTEx data browser</u>, and you can also use the requester pay google cloud bucket (gs://gtex-resources/GTEx_Analysis_v8_QTLs). Protected data, such as genotypes and metadata, were accessed via dbGaP (study accession: phs000424.v8.p2).

#### GTEx fine-mapped sQTLs and eQTLs

Baberia et al. fine-mapped sQTLs and eQTLS in GTEx using the DAP-G fine-mapping method. We accessed this dataset using their zenodo <u>link</u>. Note that fine-mapping was performed only on European Ancestry samples from GTEx.

#### gnomAD

We accessed LOEUF scores from gnomAD v4.0 using the gnomAD data browser (linked here), which computed the LOEUF across nearly all genes in the genome.

#### GENCODE exon annotations

We accessed comprehensive transcript annotations from GENCODE v44 using their data browser, for use as our reference set of exons.

#### SpliceAI scores

SpliceAI scores have been computed across the genome (all possible SNP mutations). To compare the differences in SMS and spliceAI scores, we used a dataset that released all variants with SpliceAI scores above a nominal threshold (>0.1), which are available in the illumina data browser (link: https://basespace.illumina.com/s/otSPW8hnhaZR). All other variants were set to 0.

#### Fine-mapped BioBank (GWAS) data

UKBB (96 traits): https://github.com/mkanai/finemapping-insights, including *ASGR1* exon 4

Biobank Japan: https://pheweb.jp/downloads

FinnGen: https://www.finngen.fi/en/researchers/data_available

### Data preprocessing

#### GTEx LeafCutter phenotypes

LeafCutter intron clusters were required to include only 2 or 3 introns, so we could classify them as alternative 3’ or 5’ splice sites and skipped exons, respectively. We mapped to genes and strands by scoring every splice site in the cluster, and picking the strand that had MaxEnt scores > 0 for all splice sites, excluding any clusters that did not score above zero across all splice sites in either strand. Only protein-coding genes were included. Lastly, only exons that were less than 500 bp were included

### Generating estimates of Ψ values of NVEs

Many NVEs have fairly low inclusion levels, so in order to estimate their usage accurately, we built a model to verify their presence in an RNA-sequencing sample. We considered the usage of a potential NVE in relation to the usage of a more frequently observed “cognate” junction in transcripts from the same gene.

PSI (Ψ) estimates the fraction of a gene’s transcripts that contain the exon or splice site of interest and is a widely-used statistic to quantify splicing. It can be estimated from any RNA-seq dataset with sufficient read coverage of the alternative region.

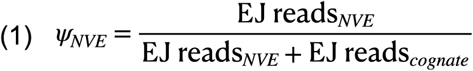

We consider the exon or splice junction with fewer reads across all individuals a candidate naturally variable exon (NVE) and the reference exon or splice junction with more reads is considered the cognate.

Conditional on Ψ, the number of NVE reads in an RNA-seq dataset is modeled as a binomial, as defined by LeafCutter splicing^54^ clusters

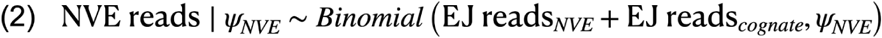

Note that we have to frame the exon junction reads slightly differently in alternative splice sites and skipped exons to model the NVE **(Extended Fig. 1A, Supplementary Note)**.

### Beta binomial model

Partial pooling involves using information across samples to estimate effect sizes of individual samples by fitting a population distribution to the observed data and letting that inform effect size estimates for individuals^55,56^. Here, we used a single-beta distribution, with parameters α and β:

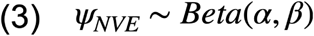

Where *α* can be described as the prior on the splicing of the NVE splicing event, and *β* can be described as the prior on the splicing of the cognate splicing event.

A single-beta distribution is able to capture several common situations (but not all – see **Supplementary Note**). The task is to predict the population frequency of a splicing event in a given tissue sample. We employ the L-BFGS-B optimization algorithm to determine the α and β parameters under which the data is most likely. The full method is represented graphically in **Fig. 1A**.

### Alternative splice site NVEs

We searched for the optimal α and β parameters that best fit the data, using the L-BFGS-B algorithm (see **Supplementary Note**), using a beta prior with parameters α = β = 1.

### Skipped exon NVEs

Skipped exons are represented by three intron clusters: two introns supporting the inclusion of the exon, and one longer intron excluding the exon. We include events where both inclusion introns have similar numbers of reads (see **Supplementary Note**), and exclude any introns for which large differences between the two introns were observed, keeping one of the two inclusion introns to simplify the analysis. We then followed the same approach as for alternative splice sites to find the optimal α and β parameters.

### Calculation of exon frequency (EF) at given Ψ threshold

We estimated the Ψ of all NVEs of length (for skipped exons) or inter-splice site distance ≤ 500 bp (see **Supplementary Note**), as this size captures the vast majority (>99%) of known internal exons and alternative splice sites in the human transcriptome. After obtaining the population parameters for all splicing events, we calculated the EF at particular Ψ thresholds by using the population distribution of Ψ across individuals in that sample. For each of four cutoff values *c* = 1%, 5%, 10%, or 20%, we computed EF as the percentage of individuals with Ψ ≥ c from the modeled CDF of the population distribution and provide this in a summary table (**Supplementary Table 1**). For a tissue with *n* individuals, we included events where the EF 5% was between 1/*n* and (*n*-1)/*n*, excluding poorly spliced exons with EF below 1/n and “canonical alternative” exons with EF > (*n*-1)/*n*.

To obtain an estimated Ψ value for an NVE in a single individual, we computed the mean posterior on Ψ and confidence intervals on the estimate in each individual in the sample by using the α and β parameters estimated from the beta binomial fit described above. The posterior mean is estimated as:

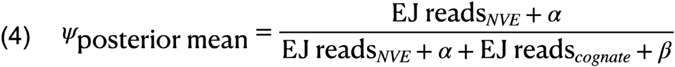

### Estimation of total NVEs per individual and number of shared NVEs between individuals

Because EF values estimated from better-sampled tissues are likely more accurate than those estimated from lowly-sampled tissues, we calculated top EF values using only the top 10 most sampled tissues from GTEx for use in estimation of the expected number of NVEs expressed in an individual. In this calculation, and the estimation of the number of NVEs shared between unrelated individuals, we assumed that NVEs occur independently of each other with probability estimated by their EF. Because other NVEs occur exclusively in tissues outside of these 10, these estimates are likely quite conservative. 57% of the data is included in the top 10 tissues, and the tissues are: Whole Blood, Thyroid, Adipose Subcutaneous, Skin (Sun Exposed Lower leg), Nerve (Tibial), Artery (Tibial), Skin (Not Sun Exposed Suprapubic), Esophagus Mucosa, Lung, and Cells (Cultured fibroblasts).

### Variant analyses

#### Fine-mapped sQTLs

As above, we filtered for intron clusters containing two introns (with one shared endpoint), which represent alternative splice sites, and clusters containing three introns in a pattern consistent with presence of a skipped exon. If the intron cluster corresponds to an NVE, then any sQTL (i.e., any variant present in the fine-mapping dataset) associated with the intron cluster is considered to be associated with the NVE.

#### Top sQTL PIP file

We created pan-tissue clusters by merging the locations of each intron in a cluster, assigning pan-tissue cluster IDs. We then sorted by PIP in descending order, and dropped duplicates based on pan-tissue cluster IDs and variant pairs, keeping the highest PIP value. This ensured that we were considering the top variant-phenotype pair for all splicing events. We performed the analogous analysis with eQTLs, except the top gene was filtered instead of cluster ID. To compute the percentage of NVEs with *cis* effects, we constructed the sQTL high confidence set: sQTLs that had a fine-mapped PIP>90%, and further filtered for variants within 5 kb of the splicing site to obtain since most splicing regulation is thought to occur within a moderate distance from the splice site^57^.

### SMS algorithm

We use fine-mapped GTEx sQTLs and filters for LeafCutter splicing phenotypes that can be categorized as exon skipping or alternative 5’ or 3’ splice sites, as above (see partial pooling section). For training, we separated out high-PIP sQTLs (90% of above) as a positive class and low-PIP sQTLs (0.2% of below) as a negative class.

For features to train the model, we used PhyloP conservation, both 5p and 3p MaxEnt splice site scores (by scanning around the variant and looking for the maximum score of each splice site), change in MaxEnt score due to variant, location within an exonic splicing enhancer^39^ or exonic splicing silencer^40^ motif, splicing-associated histone marks^41^, location within the binding site of an RNA-binding proteins, based either on eCLIP peak data^42^, or based on a mapping of in vitro-derived binding motifs in the transcriptome^58^, and the log of the distance to the nearest splice site.

We trained a logistic regression to predict feature weights using annotated feature matrix M. We found that feature weights changed depending on where the variant was present within the gene. For example, exonic splicing enhancers have stronger weights in annotated exons than in introns. The gene annotations considered are: splice sites (including either GENCODE splice sites or GTEx SS in SMS-full), GENCODE annotated exon, or GENCODE annotated intron. So, we also include a gene annotation vector A. The vector contains where the exon is in the gene.

For a given variant *i*,

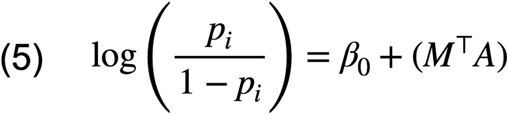

where *p_i_*is the probability that the variant *i* modifies the splicing outcome.

We trained this logistic regression model on 80% of randomly selected sQTLs to determine the relative importance of each feature in predicting causal sQTL variants.

We trained on both the full model and on subsets of the features, and computed statistics such as AUPRC of these sets of models using held-out test sets, which is reported in the text. The Github repository includes all annotations for GTEx variants and a tutorial for rerunning many combinations of features for this model.

### Splice comparison with SMS

To compare our SMS with spliceAI, we also trained a logistic regression model using SpliceAI scores downloaded from <u>basespace</u>. (Note that on basespace, all SpliceAI scores below 0.1 are listed as 0.)

### Distribution of loss-of-function intolerant genes

The loss-of-function observed/expected upper bound fraction (LOEUF) scores, which was first described in this study ^23^, were obtained from the <u>gnomAD browser</u>. We used the most recent release (v4).

### Enrichment of GWAS variants in splice sites across biobanks

First, associations of variants across all traits were concatenated into a file of trait-variant pairs. We filtered for single variant-trait pairs, taking the top PIP across all traits (so as to not repeat variants that had high PIP associations across many traits).

Then for each variant, we computed the maximum PIP across traits in BBJ, FinnGen, and UKBB, and pooled these variants together. We estimated functional enrichment for each category as a relative risk (RR, i.e., a ratio of proportion of variants) between being in an annotation and fine-mapped. That is, RR = (proportion of variants in annotation with PIP within a specific bin) / (proportion of variants in annotation with PIP ≤ 0.01).

The PIP bins we use are the following: [0, 0.01, 0.01, 0.25, 0.5, 0.75, 1.0], where all of the intermediate values are open brackets.

To obtain enrichments of GWAS variants for unannotated GTEx splice sites, we filtered out any splice sites in GENCODE, and also variants in pLoF, splice region, or, missense variants. Enrichment is calculated using a fraction of variants observed in the lowest PIP bin (see range on plot) relative to the bin in question. Enrichment for missense and synonymous changes were also computed, keeping pLoF and missense variants in that set. We provide the locations of all unannotated GTEx in **Supplementary Table 4.**

Because unannotated GTEx splice sites comprise a very small fraction of the genome and tend to occur close to existing coding regions, we did not use LDSC to assess whether unannotated GTEx contributed to SNP-heritability of traits.

### Detection of unannotated splice sites impacting NMD

We devised a script that called unannotated splice sites with NMD potential in GTEx LeafCutter splicing phenotypes. The script: 1) intersects LeafCutter introns with annotated UTRs, excluding those that overlap; 2) maps remaining LeafCutter introns to corresponding exons in annotated coding regions; 3) Assesses putative NMD potential if either a) the NVE alters the frame of an annotated CDS, e.g., adding an exon whose length is not a multiple of three, or b) the NVE contains an inframe stop codon; 4)

Excludes any NVE that impacts the last exon within the CDS; and 5) Excludes any NVE that represents the second-to-last exon of a CDS whose only stop codon(s) are within 50 bp from the NVE 5’SS. This script is intended to capture common NMD-triggering splicing changes, but does not cover some special cases. For example, if a pair of adjacent NVEs in the same gene each individually preserve frame they will be considered frame-preserving, even if when spliced together they generate a stop codon at the exon-exon junction. Conversely, if two NVEs each alter the reading frame, they will be considered as having NMD potential even if when they are spliced together the reading frame is restored.

### sQTL that we also eQTLs

First, GTEx variants were filtered for those that were both nominally significant sQTLs (.v8.sqtl_signifpairs.txt) and nominally significant eQTLs (.v8.egenes.txt) in the same tissue <u>(GTEx browser)</u>. The sQTL phenotypes considered only included splicing events that were alternative splice sites, where one of the sites was an NVE. The exact sQTL phenotype was the cognate intron, and not the NVE, because not many variants were nominally significant in the other direction (impacting the NVE levels), likely due to lower Ψ values of NVEs, so we transformed the effect size by -1 for all sQTLs, in order to more directly compare the impact of the NVE inclusion on expression of the gene.

### Data availability

All data is publicly available. Tables of EF_5%_ values are included in a zenodo bucket.

### Code availability

<u>Code</u> for calculating the alpha and beta parameters of NVEs called partial pooling variable splicing (PoVS) is posted as a docker image on DockerHub. <u>Code</u> for a tutorial for running this docker image interactively is on Github, including some test data from GTEx. The rest of the data can easily be accessed in the <u>GTEx data browser</u> to run all other files and generate the EFs for all tissues.

Code used to post-process these events (called post-process-PoVS) is on Github, and can be used to generate the EFs of NVEs reported in **Supplementary Tables 1 & 2**.

Code for logistic regression is also available on Github called splice modifier score (called SMS), including a jupyter notebook which can be used to retrain the model using different features, or regenerating SMS full model predictions (**Supplementary Table 3**).

### Additional Information

More detailed methods are presented in the **Supplementary Note**, which include figures to explain the underlying model behind the partial pooling method and QC steps for NVEs.

## Supplementary Tables

Supplementary Table 1 maxent filtered, size filtered, protein coding NVEs across all tissues in GTEx

Supplementary Table 2 maxent filtered, size filtered, protein coding NVE in tissue with highest frequency (top EF) in GTEx

Supplementary Table 3 SMS scores for all GTEx variants, also annotated with spliceAI scores

Supplementary Table 4 Splice site regions present in GTEx and not GENCODE used for fine-mapping enrichments

## Extended Data

**Extended Data Fig. 1.**
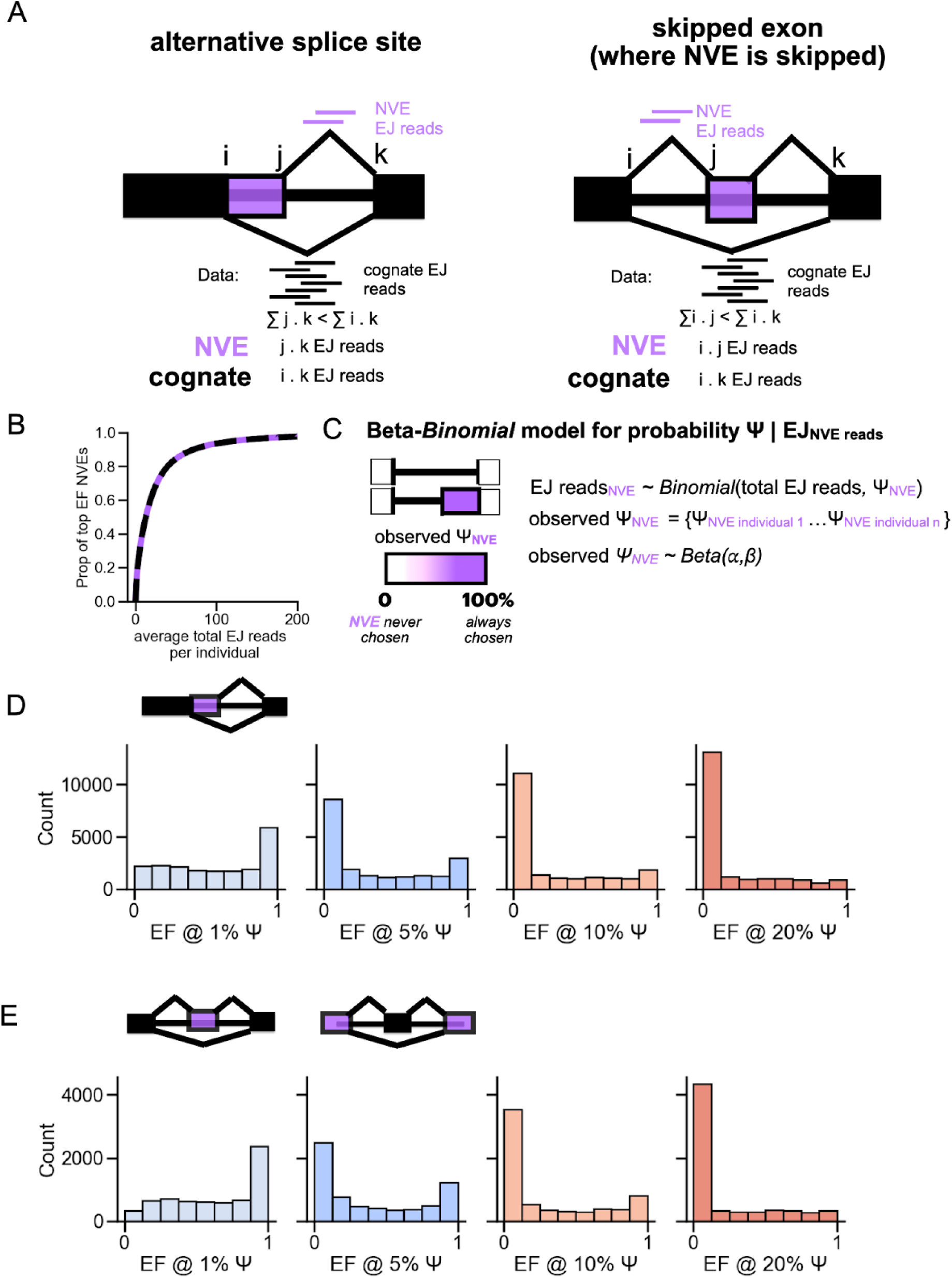
Beta-binomial method and EF distributions at different PSI cutoffs A) Illustration of how the EJ reads are chosen to be put into the beta binomial model for fitting. Note on the right that the NVE is skipped, but NVE can also be not skipped - the general logic is the same. B) Mean total EJ reads (cognate + NVE) per individual for top EF NVEs. Computed using the total EJ reads (cognate + NVE) divided by the number of people in the tissue. C) Explanation of the beta binomial model applied to EJ count data. D) EF spectrum of alternative splice site NVEs at different minimum Ψ thresholds, for one tissue (Heart Atrial Appendage) E) EF spectrum of skipped exon NVEs at different Ψ thresholds, for one tissue (Heart Atrial Appendage).

**Extended data Fig. 2.**
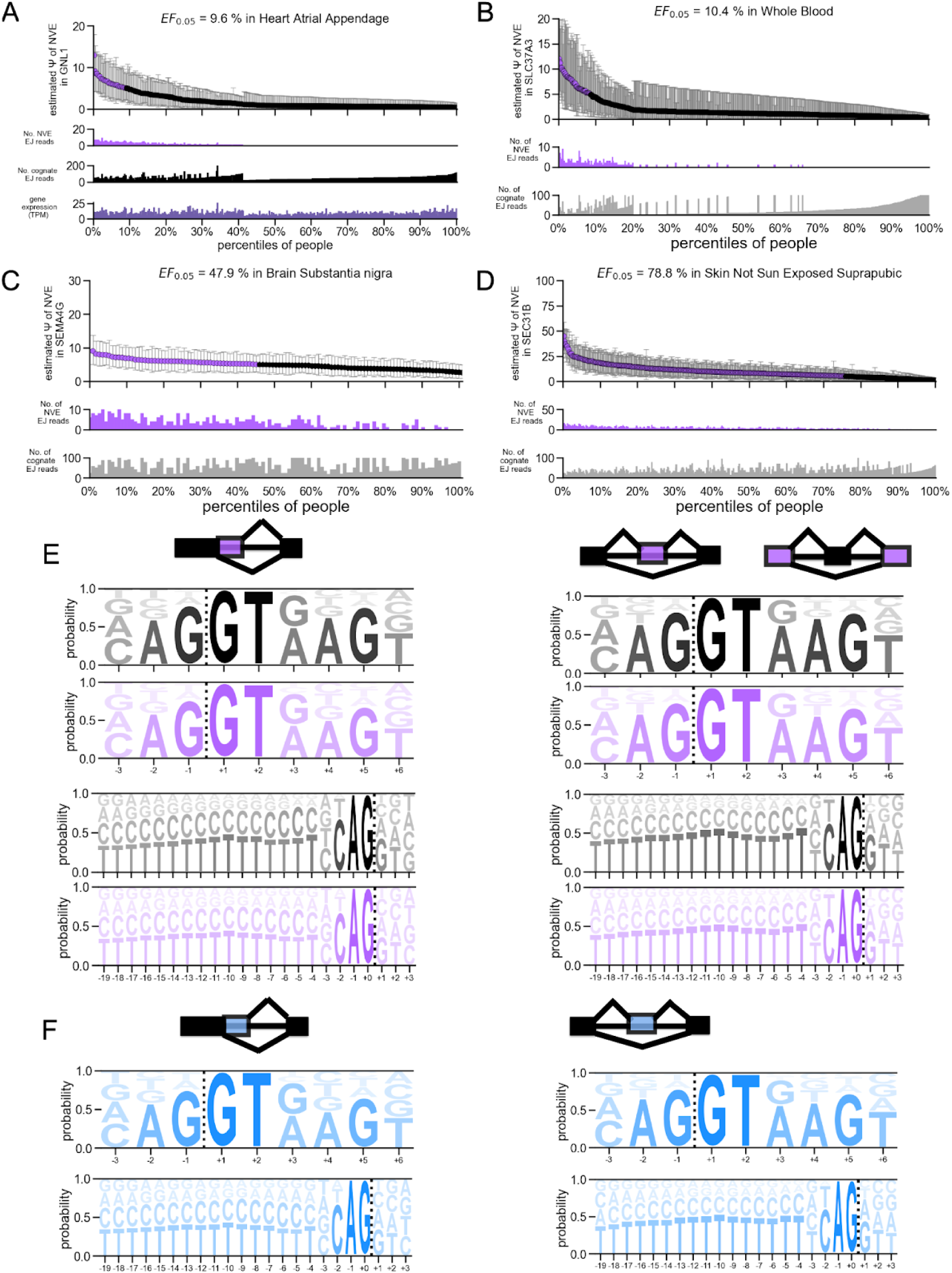
Single exon plots showing dynamic nature of NVEs across EFs and splice site motifs of cognate and NVEs. A) Detailed exon locus plot for an NVE in *GLN1*, highlighted in main text (Fig. 1B). The number of reads for the NVE is in purple, shown as a barplot, and the number of reads for the cognate exon is in gray. TPM of *GLN1* is shown in dark purple. Posterior mean estimates (PME) of Ψ for an exon is plotted (dots), along with 95% credible intervals on the estimate (error bars). The PME is colored purple if it meets the threshold of 5% Ψ in the individuals, black if it does not cross that threshold. The EF_5%_ is plotted on top, which, definitionally, should generally mirror the proportion of exons in purple vs black. B-D) Exon locus plots at different EF _Ψ 5%_ values. The number of reads for the NVE is in purple, shown as a barplot, and the number of reads for the cognate exon is in gray. Posterior mean estimates (PME) of Ψ for an exon is plotted (dots), along with 95% credible intervals on the estimate (error bars). The PME is colored purple if it meets the threshold of 5% Ψ in the individuals, black if it does not cross that threshold. The EF_5%_ is plotted on top, which, definitionally, should generally mirror the proportion of exons in purple vs black. Note the scales for barplot differ based on the distribution of reads for each exon. E) Left) Splice site motifs across alternative splice sites. NVE shown in purple, cognate shown in black. Right) Splice site motifs across skipped exons. NVE shown in purple, cognate shown in black. F) Splice site motifs across low EF alternative splice sites, shown in light blue (to represent its low EF value).

**Extended Data Fig. 3.**
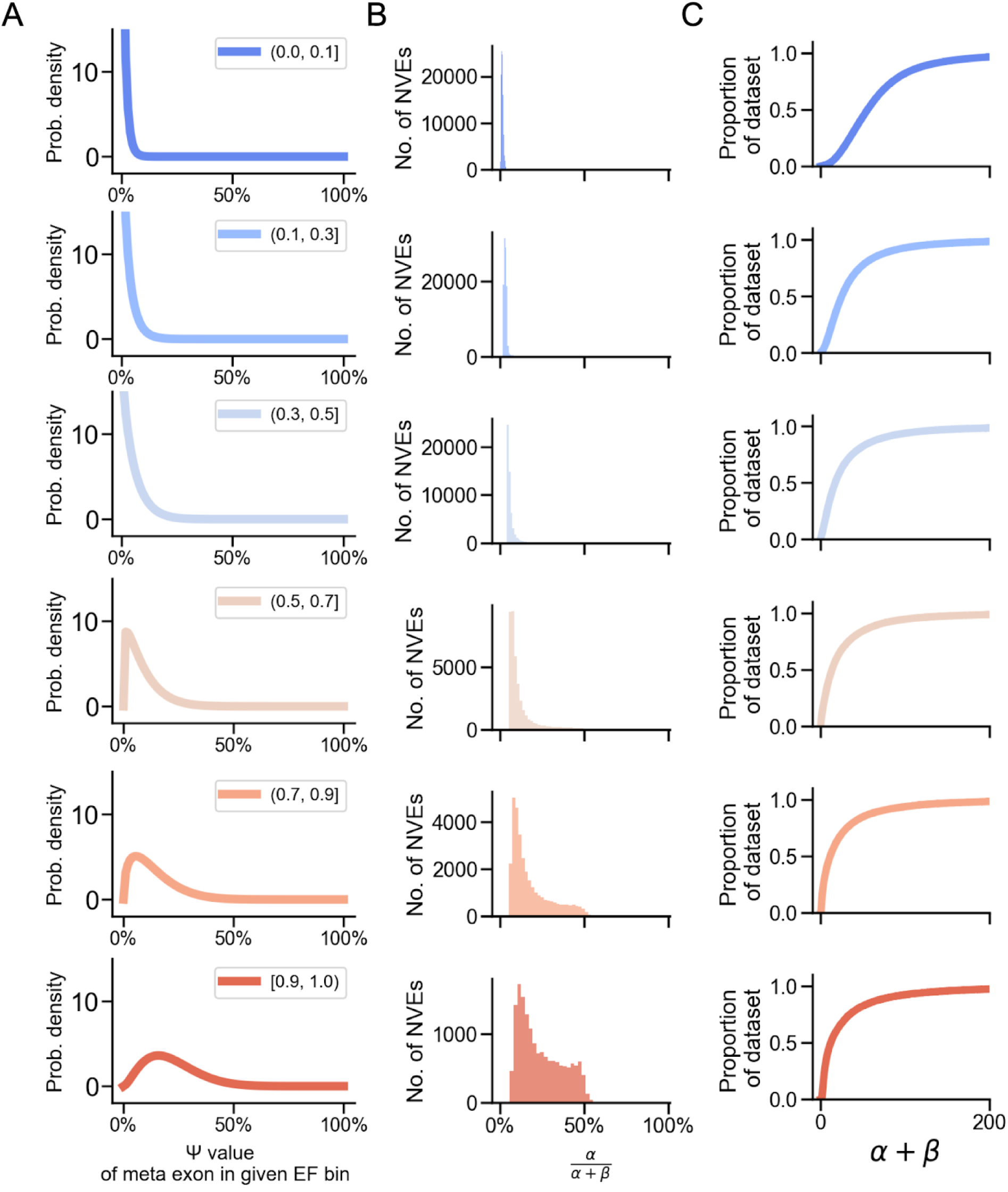
NVE summaries using population parameters. A) the median α and β values in each EF bin was computed, and then a “meta exon” was constructed using those values. Plotted is the PDF across given Ψ values for the meta exon given the specified α and β parameters. B) For all NVEs across all tissues, NVEs are separated by EF bin, and the mean estimate of Ψ is plotted. C) Same as A, except the sum of alpha and beta is plotted, which is equivalent to 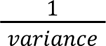 of the β distribution.

**Extended Data Fig. 4.**
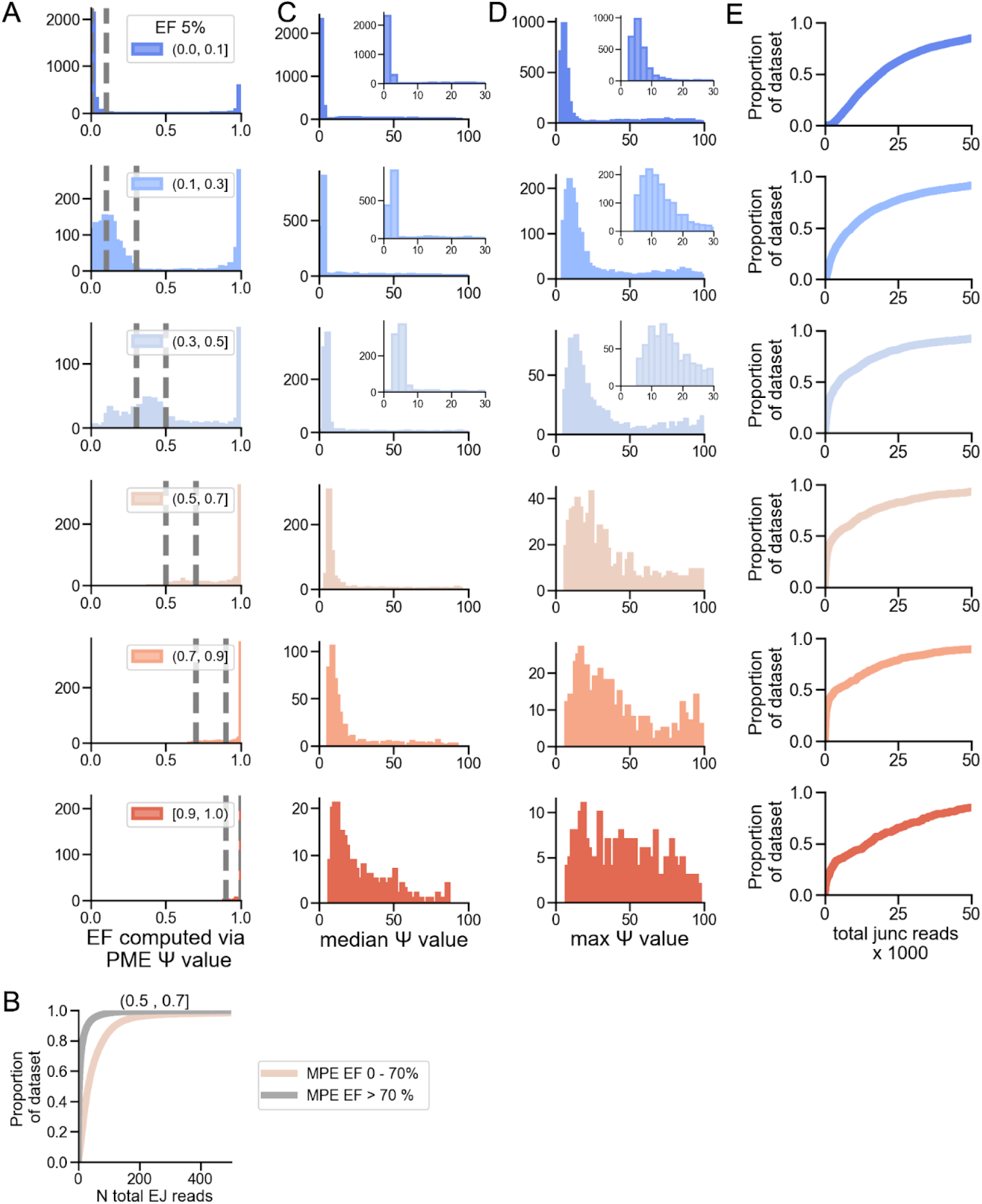
NVE summaries estimated using observed data in conjunction with population parameters. A) Relationship between EF calculated using the population distribution (typical way) and the EF calculated using posterior mean estimate (PME) on Ψ. First, a single tissue was chosen (adrenal gland), and exons were filtered for alternative splice site NVEs. EF using PME was calculated as follows: determining the fraction of the individuals with the mean posterior estimate on Ψ≥5% in that tissue. Gray lines represent the EF boundaries calculated via the first method. B) Explanation of non-concordance between two methods to calculate EF. Exons are split by whether there is concordance (tan) or not (gray) between the two ways to calculate EF. Then, the number of reads associated with both the cognate and NVEs are plotted. Nonconcordant exons tend to have much lower reads, suggesting that using the PME estimate may not be sufficient in cases with low read depth, demonstrating the utility of using EF derived from population parameters. C) Distribution of median Ψ across different EF_5%_ frequencies. A tissue at random (adrenal gland) was chosen to generate these plots. The PMEs were determined for all exons. The median PME Ψ value across all individuals in the exon is plotted as histograms. Exons are split by their overall exon frequency in the tissue. First three EF bins contain insets to provide a more detailed view of the distribution, since median Ψ values are low. D) distribution of maximum Ψ across different EF_5%_ frequencies. First, the PME of Ψ was determined for all exons. The maximum PME Ψ value across all individuals in the exon is plotted as histograms. Exons are split by their overall exon frequency in the tissue. First three EF bins contain insets to provide a more detailed view of the distribution, since median Ψ values are low. E) The total exon junction reads across cognate and NVE, for exons across the EF_5%_ spectrum. The sum of the exon junction reads was determined for all exons across all individuals for that tissue, and then plotted via their EF_5%_ frequency bins.

**Extended data Fig. 5.**
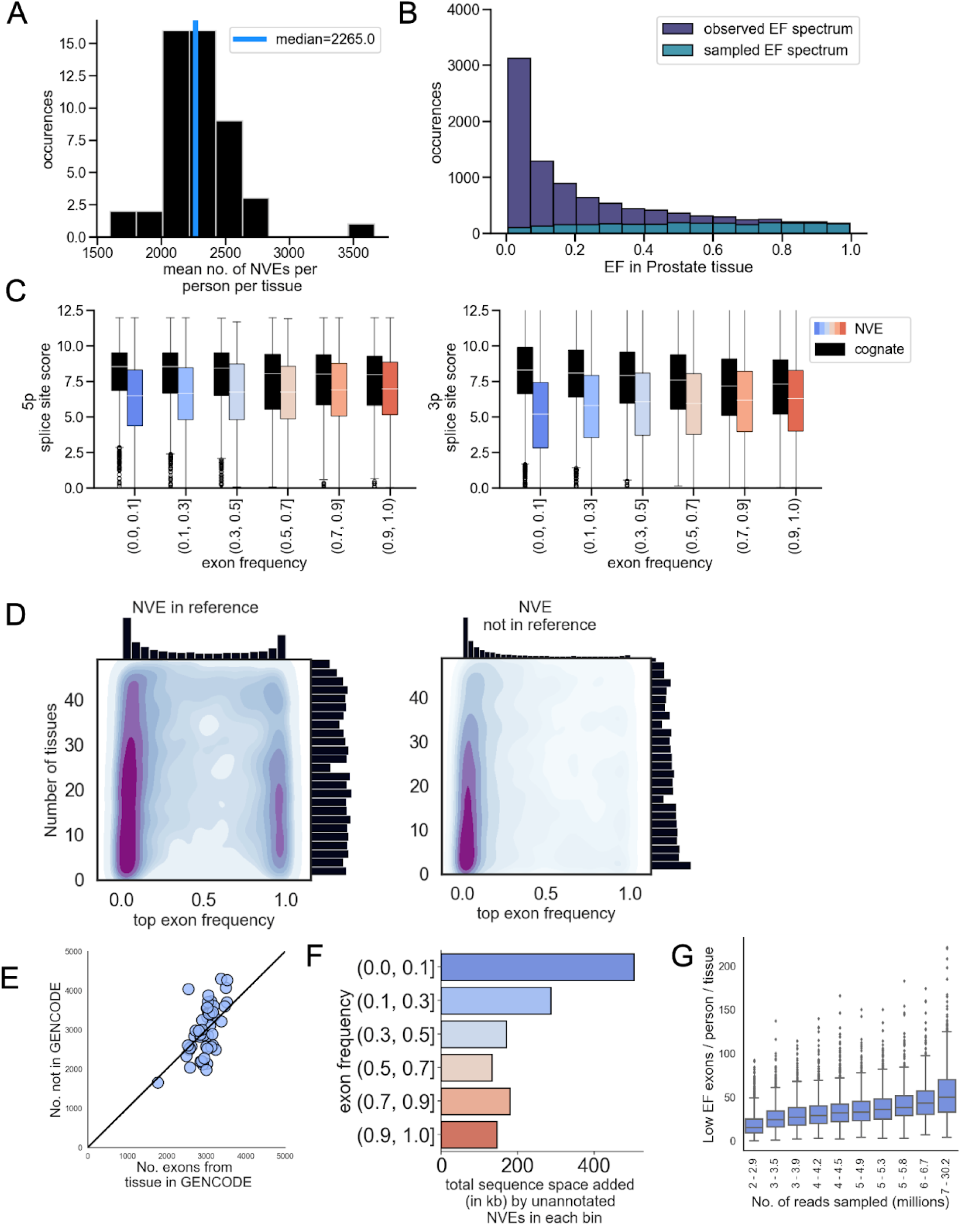
Validation and characterization of NVEs A) Median number of NVEs in tissues, across all tissues in GTEx. First, the number of NVEs per person was estimated across each tissue by taking the sum of the EFs. Then plotted as a histogram, with median NVEs/tissue plotted. B) First, EFs in an individual tissue were selected (prostate). Random variables were selected in the Bernoulli trial using their EF as the probability of observing the event, done for all NVEs in the tissue (using the NVE EF as the probability). The distribution of EFs conditioned on the exon being observed in an individual, called the sampled EF spectrum, is then plotted in teal. For reference, the dark blue overall EF spectrum for Prostate tissue is plotted. C) Maxent splice site motifs scores for 5’ NVE_alt ss_ (left) and 3’ NVE_alt ss_ (right). Cognate shown in black and NVE shown by their EF bin. D) Density heatmap is computed by counting up the number of times an NVE is seen in a tissue in GTEx, and plotting that value vs top EF. In the left plot, only NVEs in the reference database are included, and you can see the majority are pan tissue low EF exons or high EF exons. The right plot is only showing non-reference NVEs. Majority of these NVEs are low in EF and number of tissues seen. E) showing the number of exons that are in reference and non reference sets in each tissue as dots, with 1:1 ratio line plotted for scale. F) additionally coding space occupied by non-reference NVEs in the transcriptome, binned by EF frequency G) relationship between read depth of sample and number of low EF NVEs detected in the sample.

**Extended Data Fig. 6.**
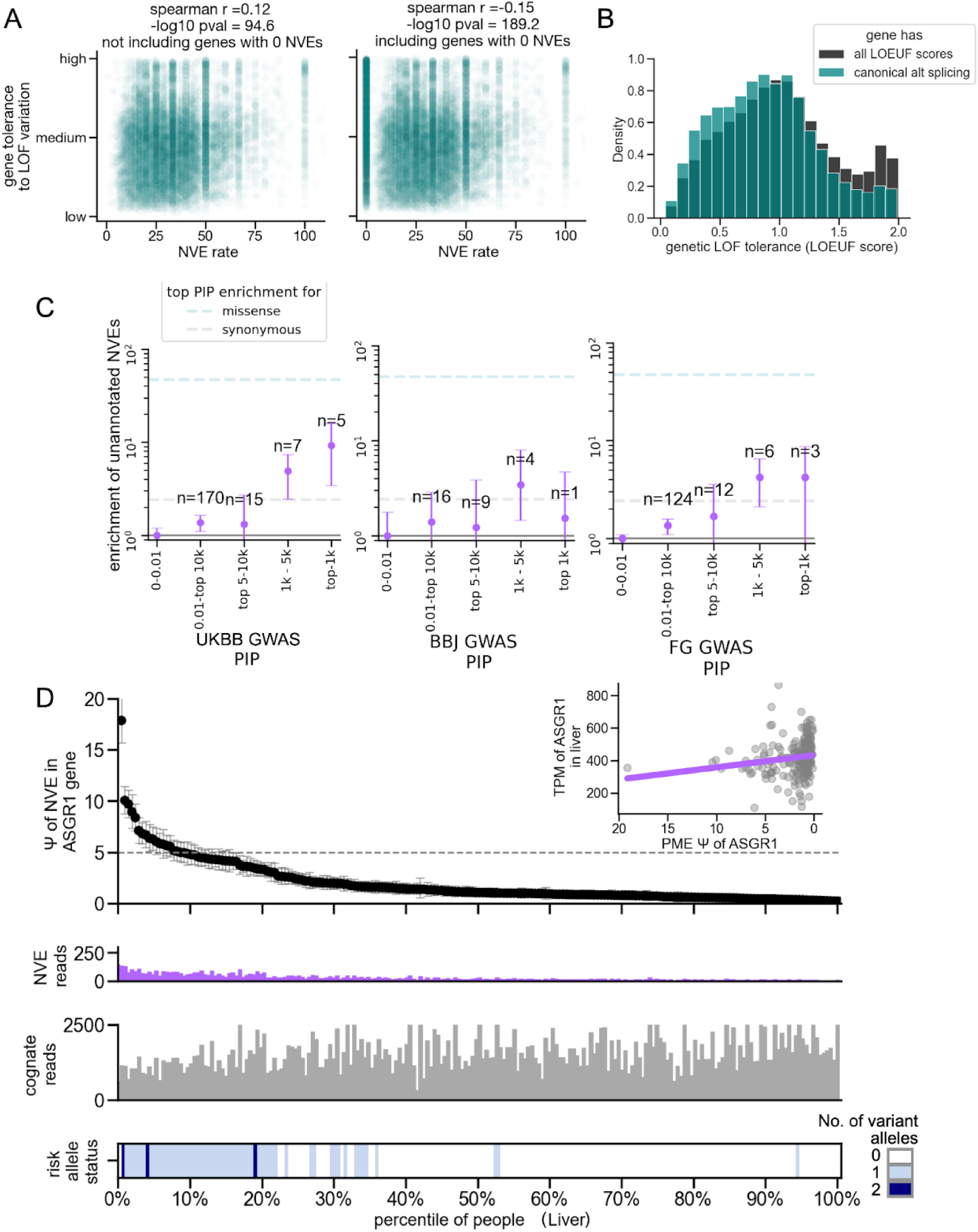
NVE LOEUF score analysis and predictive power of NVEs across cohorts A) Spearman correlation between NVE rate (number of NVEs per gene divided by the number of exons in reference transcriptome, highly significant: -log_10_pval > 90) and the LOEUF score of the gene. Right hand plot shows correlation with genes with 0 NVEs added, right hand plot shows correlation without them. B) LOEUF scores of all genes (background), vs GTEx alternative splice sites and skipped exons that were not NVEs (instead, they were included in all individuals in the tissue) C) Enrichments of GWAS variants for unannotated splice sites, filtered out for variants in pLOF, splice region, or, missense variants. Enrichment is calculated using fraction of variants observed in lowest PIP bin (see range on plot) vs the bin in question. Due to power differences, bins were chosen at higher end by variant rank. Credible intervals were generated using 2.5-97.5 percentile bounds on binomial distribution given no. of observations. Colored lines indicate the missense and synonymous enrichment point estimates for the highest variant rank set. Number of variants being categorized as within an NVE is shown as n values. D) Synonymous GWAS variant in *ASGR1*, located within an NVE (7% EF) splice site boundary, and is a positive sQTL (presence of variant induces splicing change) in liver and also associated with liver, heart, and blood phenotypes in GWAS. Inset plot shows the correlation between TPM and PSI of exon, which has the expected negative correlation, as the GWAS variant is both a negative eQTL and predicted to cause NMD.

**Extended Data Fig. 7.**
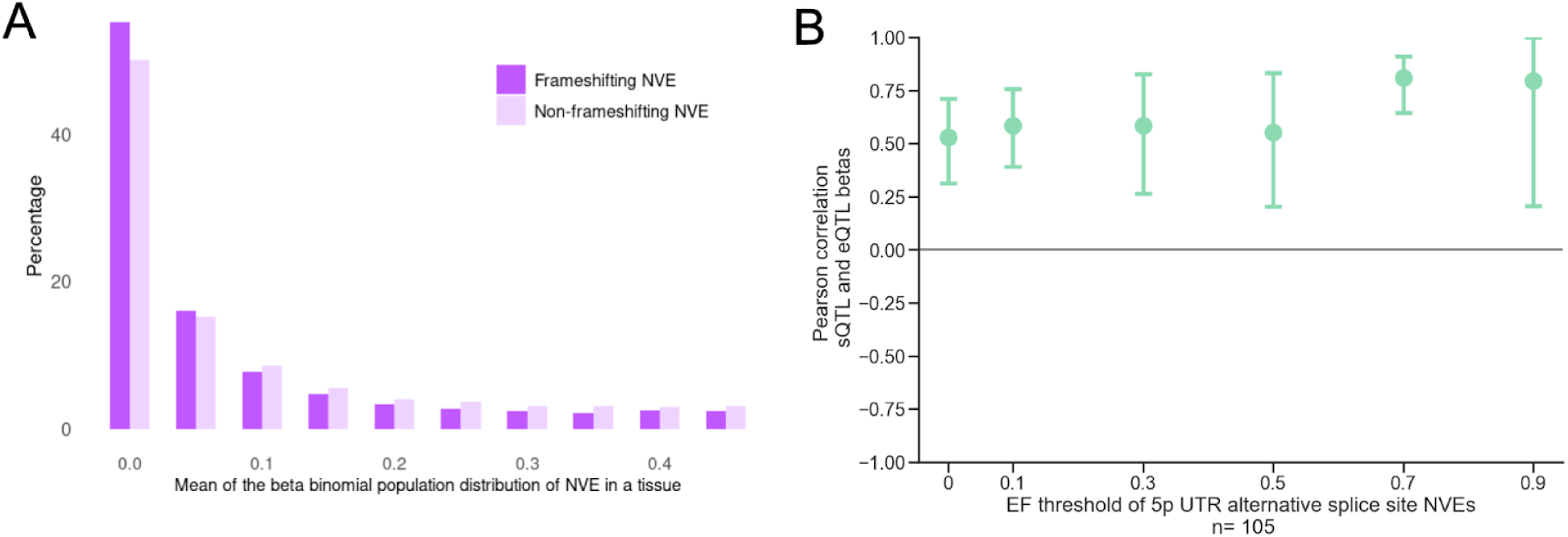
Investigation of NVEs that impact expression A) α and β parameters derived from the fitted beta population distribution were used to generate the mean estimate of Ψ for a given NVE within a given exon, and then split between NMD-potential (darker purple) and non-NMD-potential exons (lighter purple). B) NVEs that intersected within 5’UTR regions were selected. Effect size estimates (slope of regression of QTL) shown for genetic variants that were nominally significant sQTLs for 5UTR NVEs, that are also significant eQTLs in the same gene in the same tissue. Genetic variants were filtered out if in complex loci (ie, were eQTLs in other genes in the tissue). Note that association between variant and splicing event is with the cognate splicing event, such that an decrease in splicing would lead to an increase in the overall NVE:cognate ratio, so we transformed this effect size by multiplying by -1. Shown is 1000 bootstrapped Pearson correlations between sQTL and eQTL effect sizes, performed at increasing EF thresholds. The sample size for 3p UTRs was too small to analyze (23 variants).

**Extended Data Fig. 8.**
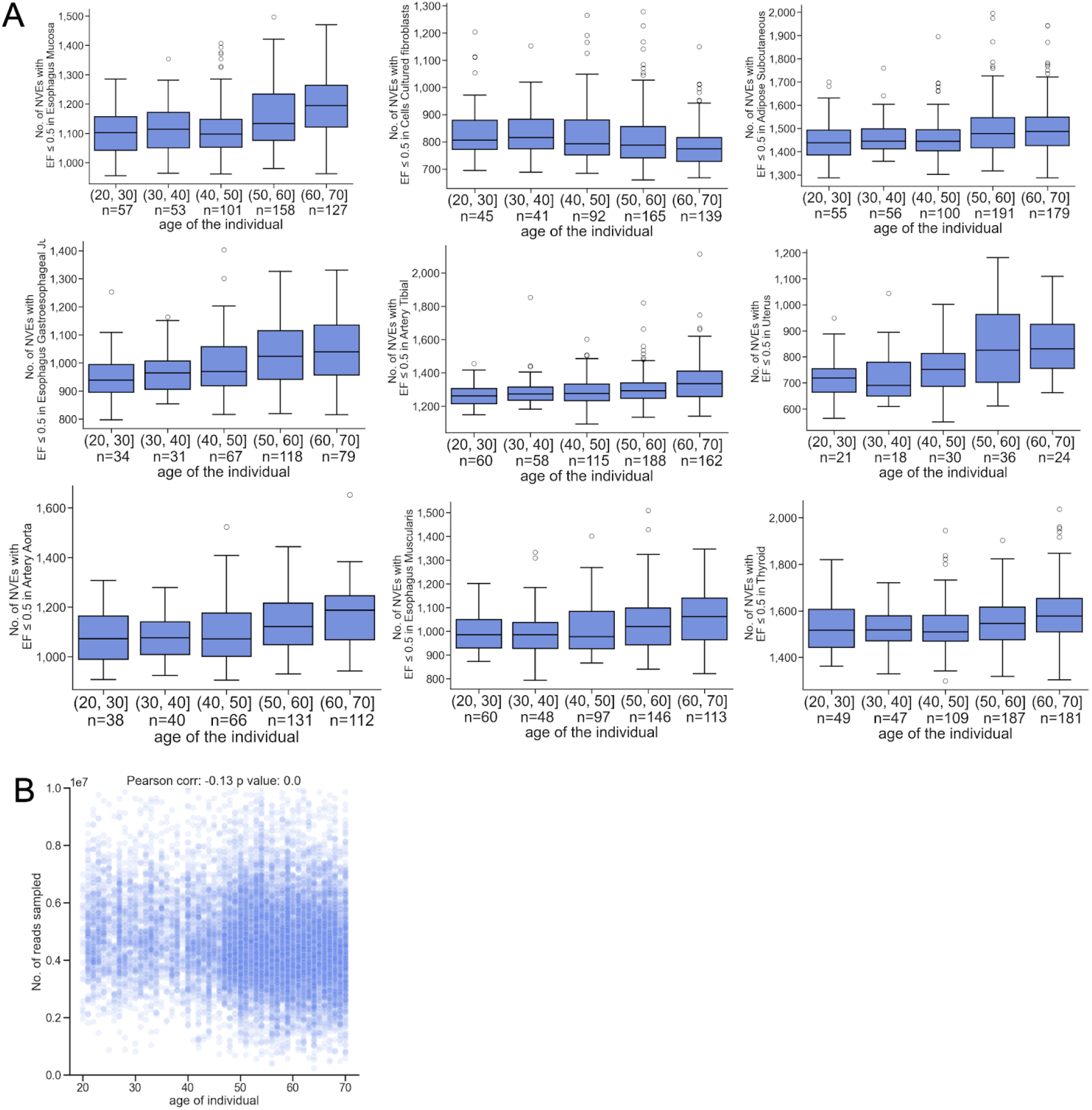
Age is positively associated with the occurrence of NVEs. A) Tissues with selected marginally significant (BH corrected alpha 0.05) spearman correlations between age and low-to-medium EF (≤50%) NVEs. y-axis is the number of NVEs vs. age of an individual on x. Each dot is an individual in the tissue sample. Note that the y-axis differs among plots. B) Pearson correlation between age of individual and no of reads in the sample.

**Extended Data Fig. 9.**
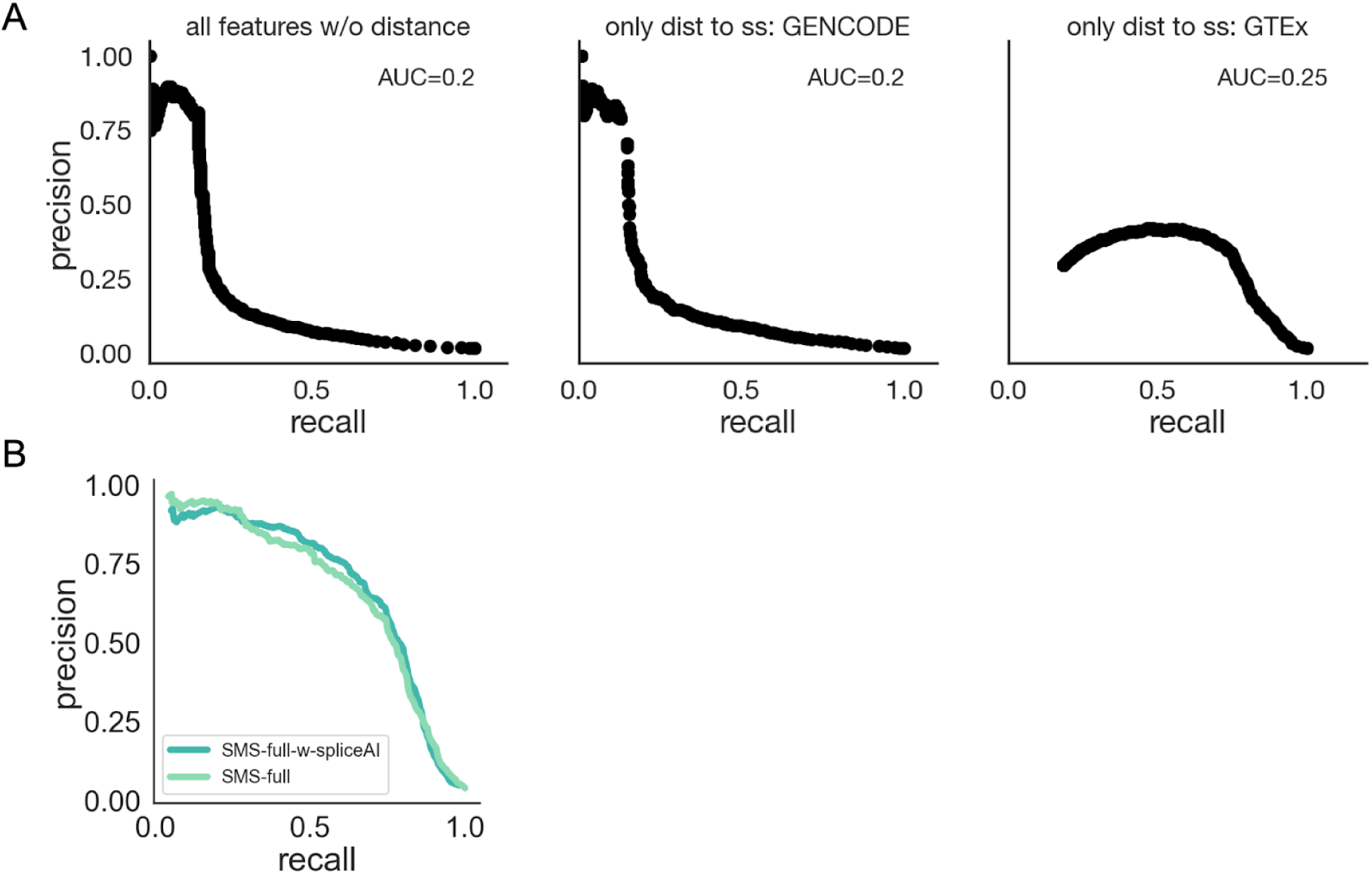
Model performance with varying features A) Performance of model in held-out test sets, using either all features without distance to splice site (far left), or only distance to splice sites as features: using GENCODE splice sites (middle) or using distance to GTEx splice site as a single feature model (right). Model was tested using a held out test set of 20% of negative and positive set variants AUC shown on top of all plots for respective models). B) Performance comparison between SMS-full (light mint) and SMS-full + spliceAI predictions (dark mint).

## Supplementary Note

<u>A. Beta-Binomial model to describe splicing A1i. Interpreting k</u>

<u>A1ii. Defining k for skipped exons</u>

<u>A2. Single beta distribution for splicing</u>

<u>B. The Objective function for the beta distribution</u>

<u>B1. Parameter settings and transformations</u>

<u>C. Choice of population distribution for EF calculation</u>

<u>D. Filtering LeafCutter outputs</u>

<u>E. Choice to not use LD score regression to assess NVE annotations</u>

### A. Beta-Binomial model to describe splicing

First, we assume that in each individual the number of reads supporting the inclusion - versus exclusion isoforms follow a binomial distribution.

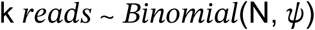

k is exon junction reads spanning the NVE

N is exon junction reads spanning the NVE and cognate

*Ψ* is the percent spliced in of the NVE (so that the cognate PSI is 1 – *Ψ*)

#### A1i. Interpreting k

The two choices in the binomial model are: NVE (k reads) or cognate (N–k reads). For exon skipping, spliced-in can either be an inclusion event or exclusion event of a given exon, and for alternative splice sites, spliced-in is the inclusion event of a particular splice site.

##### Defining naturally variable splicing events (NVEs)

We empirically define the candidate “NVE” as the less used exon, by considering the exon that has less reads in total across all individuals. In this dataset, we only perform the analysis on exons that could fall under the categories of skipped exons or alternative splicing events (see **Methods)**, both of which can be framed under our model’s single beta binomial framework.

A1ii. Defining k for skipped exons

##### Skipped exon case

There are three sets of reads that can serve as k in the case of skipped exons:

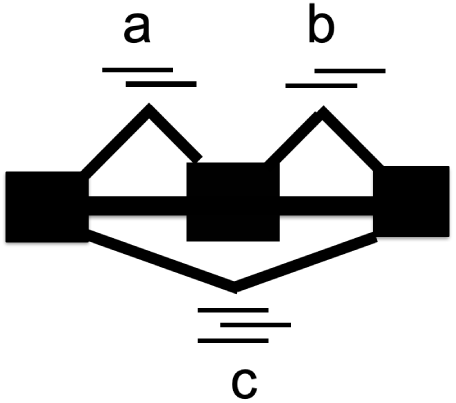

Below, we show the ratio of mean of the log counts of sets a and b for given exons in a single tissue. Though they should, at high read depth, follow an approximately 1:1 ratio if they are indeed all skipped exons. As a threshold, we require an absolute log 10 ratio less than 1, a threshold which retains the bulk of the data, including many exons with somewhat imbalanced ratios resulting from low read counts. Excluded are some extremely biased cases that likely represent complex splicing events (for example, a skipped exon and alternative splice site, or multiple alternative splice sites), which we wish to exclude. After this filter, we use the first intron present in the transcriptome to represent the reads supporting the skipped exon.

**Supplementary Figure 1.**
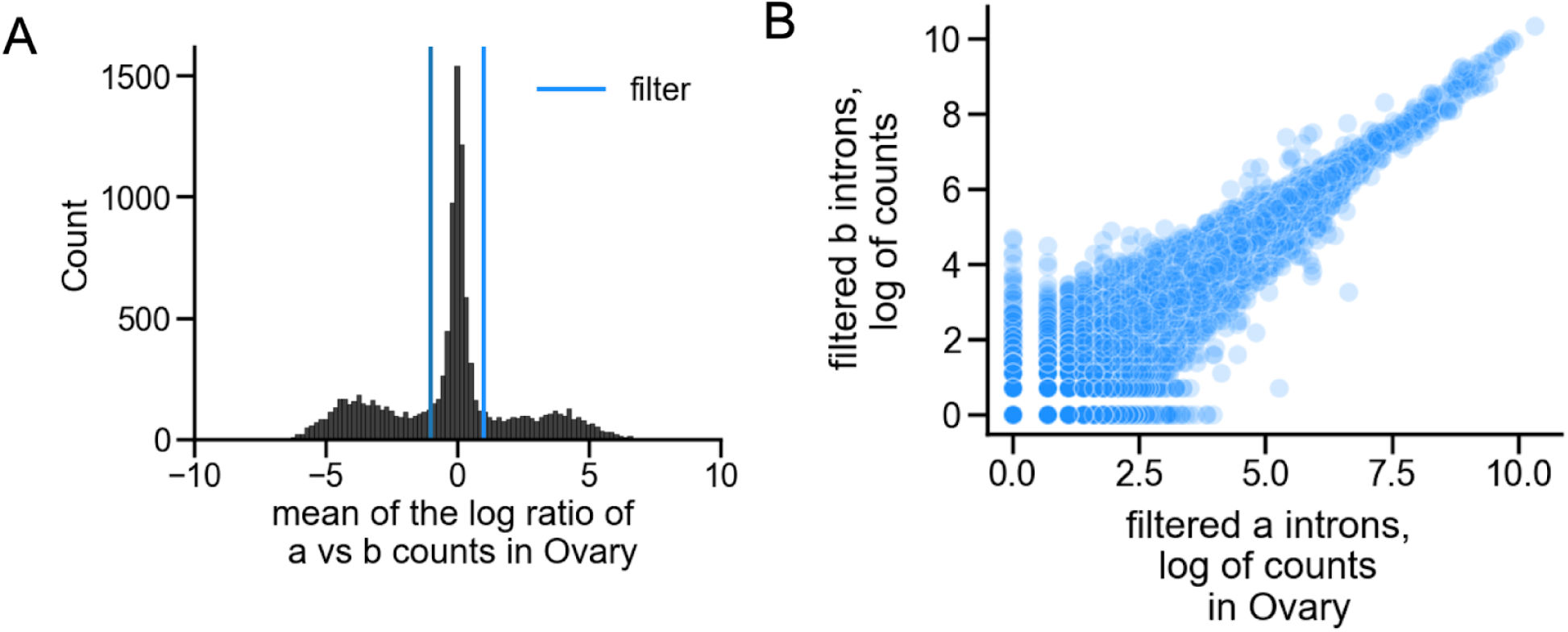
A) comparison of the mean of the log of counts for each unskipped junction of skipped exons (a and b referred to above) in an arbitrarily selected tissue: Ovary. Blue line indicates the range around 0 that passed our filter (absolute mean log ratio < 1). B) The read counts of introns a and b for NVEs that passed filtering in arbitrarily selected tissue, Ovary (Pearson and spearman correlation 0.97, 0.95 respectively).

### A2. Single beta distribution for splicing

#### Choice of single beta distribution to model splicing

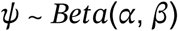

We model PSI as a beta distribution, with parameters alpha (α) and beta (β) which can capture the following types of distributions (see Table 1):

1) The density is concentrated around a fixed value, which could be 0% (rarely/ never spliced in), 100% (always/almost always spliced in), or any value in-between.
2) Uniform distribution
3) U-shaped distribution (tendency for NVE to be either lowly included or highly included

**Supplementary Table 1.**
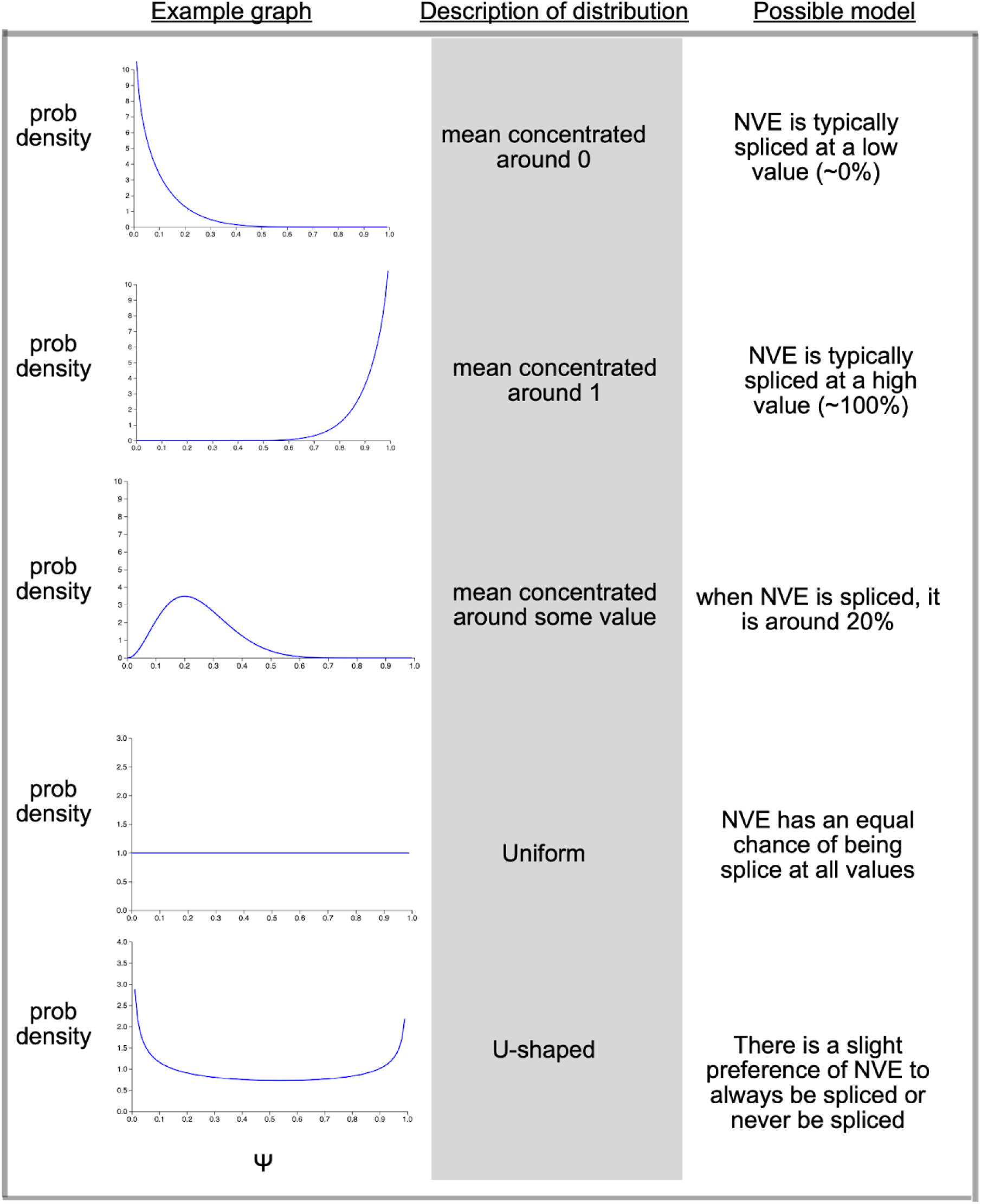
Possible modes of splicing within different beta distributions. Graphs were generated using Eureka Statistics PDF grapher.

| Example graph | Description of distribution | Possible model |
| --- | --- | --- |
|  | mean concentrated around 0 | NVE is typically spliced at a low value (~0%) |
|  | mean concentrated around 1 | NVE is typically spliced at a high value (~100%) |
|  | mean concentrated around some value | when NVE is spliced, it is around 20% |
|  | Uniform | NVE has an equal chance of being splice at all values |
|  | U-shaped | There is a slight preference of NVE to always be spliced or never be spliced |

### B. The Objective function for the beta distribution

#### The objective function

We take the sum of the log likelihood of observing the data given a set of α and β parameters with a normally distributed prior, for all values of NVE junction reads (k) and total reads (NVE + cognate junctions) (n) in individuals in the sample. This is the partial pooling step because it considers all individuals to compute the log likelihood.

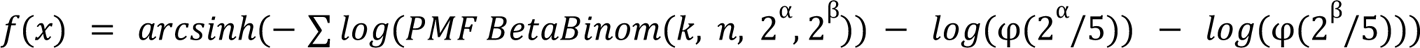

We employ the L-BFGS-B optimization algorithm to minimize the objective function.

### B1. Parameter settings and transformations Initialization

We start the optimizer with parameters α = β = 1, which is a beta distribution where all Ψ values have the same probability.

#### Including priors on α and β

We included a prior on α and β. The log2 of these parameters is assumed to follow a normal distribution with mean 0 and standard deviation of 5.

#### Exponential transformation

An exponential transformation guarantees that the parameters of the beta distribution are positive. The exponential transformation improved the numerical performance compared to using box constraints in the optimization algorithm in our numerical experiments on one tissue, both in terms of speed (roughly twice as fast) and objective value (similar for most sites, up to 20% better in some sites). The exponential transformation is reflected in the objective function above.

#### Inverse hyperbolic sine transformation

We have added an inverse hyperbolic sine transformation because it provided an additional runtime reduction of 10-20% while slightly improving the objective value in our numerical experiments on one tissue.

We provide a script that compares the exponential transformation and inverse hyperbolic sine transformation on performance and runtime (**Code availability)**.

### C. Choice of population distribution for EF calculation

Here we briefly outline our approach to calculating exon frequency. For each NVE, we use the cumulative distribution function of the beta-binomial distribution to compute the fraction of the population whose ψ exceeds the desired threshold (typically 5%). An alternative approach would be to calculate the mean posterior estimate for estimates of ψ across all individuals for the NVE, and then calculate the proportion of individuals whose ψ exceeds the threshold. However, this approach may become unstable at low read depth, since the uncertainty in ψ estimates becomes larger.

### D. Filtering LeafCutter outputs

We used all alternative splice sites and skipped exon as inputs for maximizing the likelihood function. Then, we applied the following heuristics to filter these outputs.

We used 500 bp as a filter for the size of splicing events. Specifically, we only considered skipped exons or alternative splice sites that were at most 500 bp in size. We show that there is no appreciable difference in distribution of skipped exon size, even if the NVE is a skipped exon (**Supplementary Note Fig. 2B**). As a sanity check, we provide detailed summaries of the proportion of NVEs in GENCODE annotations across all categories of alternative splice sites and skipped exons (**Supplementary Note Fig. 2A**), demonstrating that most introns are present in annotations, with the notable exception of NVE introns of alternative splice sites.

The LeafCutter outputs did not have a strand assigned. We used <u>MaxEnt</u> scores of splice sites to assign a strand to the splicing events detected by LeafCutter. MaxEnt yields a score that represents the log-odds that the site in question is a 5’ or 3’ splice site rather than a background transcript position. The scores of splice sites for constant regions, which are the regions that are shared by both splice sites, are compared with the scores of cognate and NVE exons for alternative splice sites (**Supplementary Fig. 3**) and skipped exons (**Supplementary Fig. 4**). We compare MaxEnt scores across the EF spectrum in **Supplementary Note Fig. 5**. These figures show that as EF increases, the MaxEnt scores between the cognate and NVE splicing events converge. The comparison of MaxEnt scores across the EF spectrum for alternative splice sites is shown in **Extended Data Fig. 5A**.

**Supplementary Figure 2.**
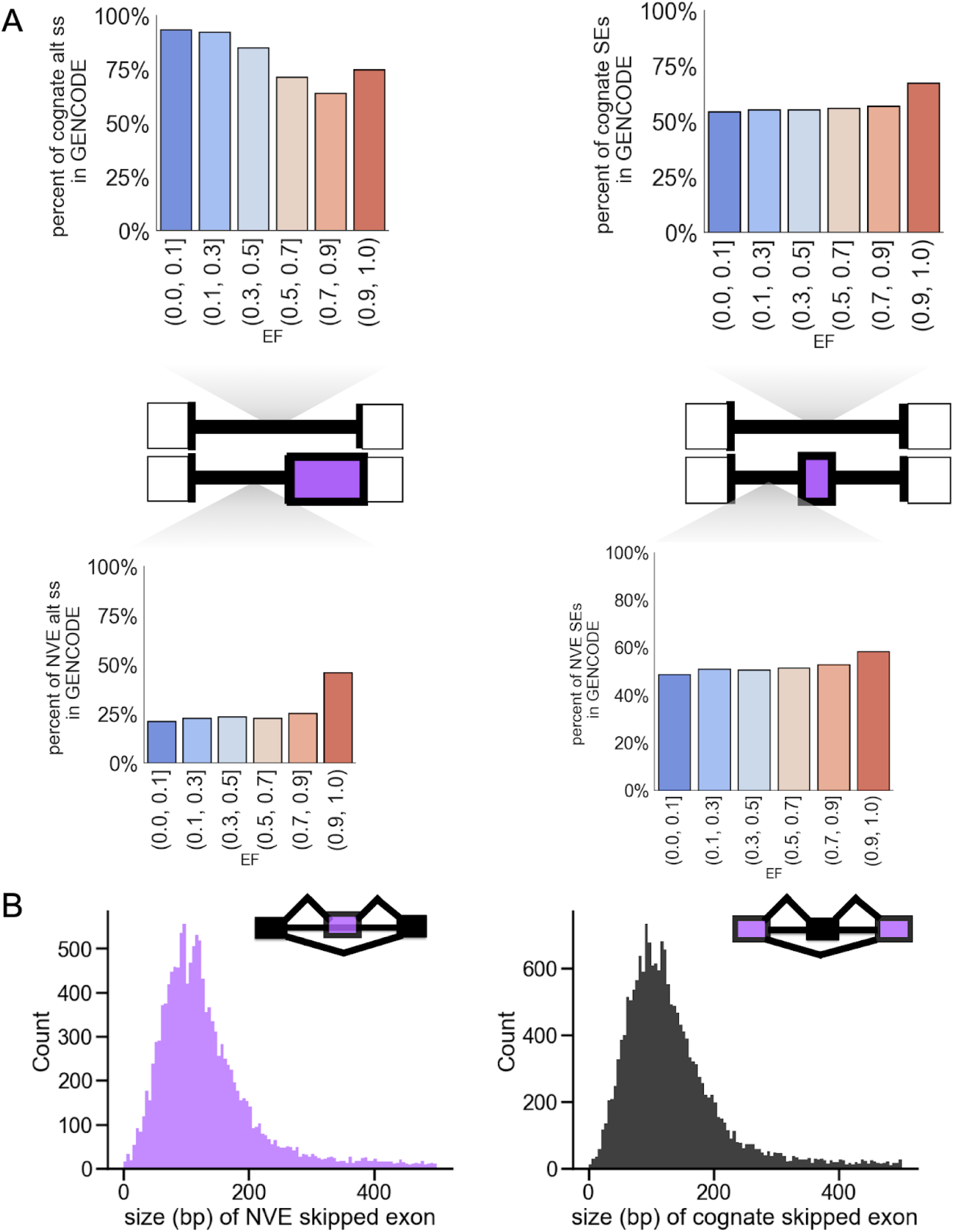
GENCODE reference intersection of NVEs, and size distribution for different classes of skipped exons. A) Fractionof cognate (top plots) and NVE (bottom plot) introns that fall within GENCODE regions are plotted, and conditioned on EF bin. The statistics are stratified by alternative splice sites (left) and skipped exons (right). B) Size in bp of each skipped exon, depending if the skipped exon is NVE (left) or cognate (right).

**Supplementary Figure 3.**
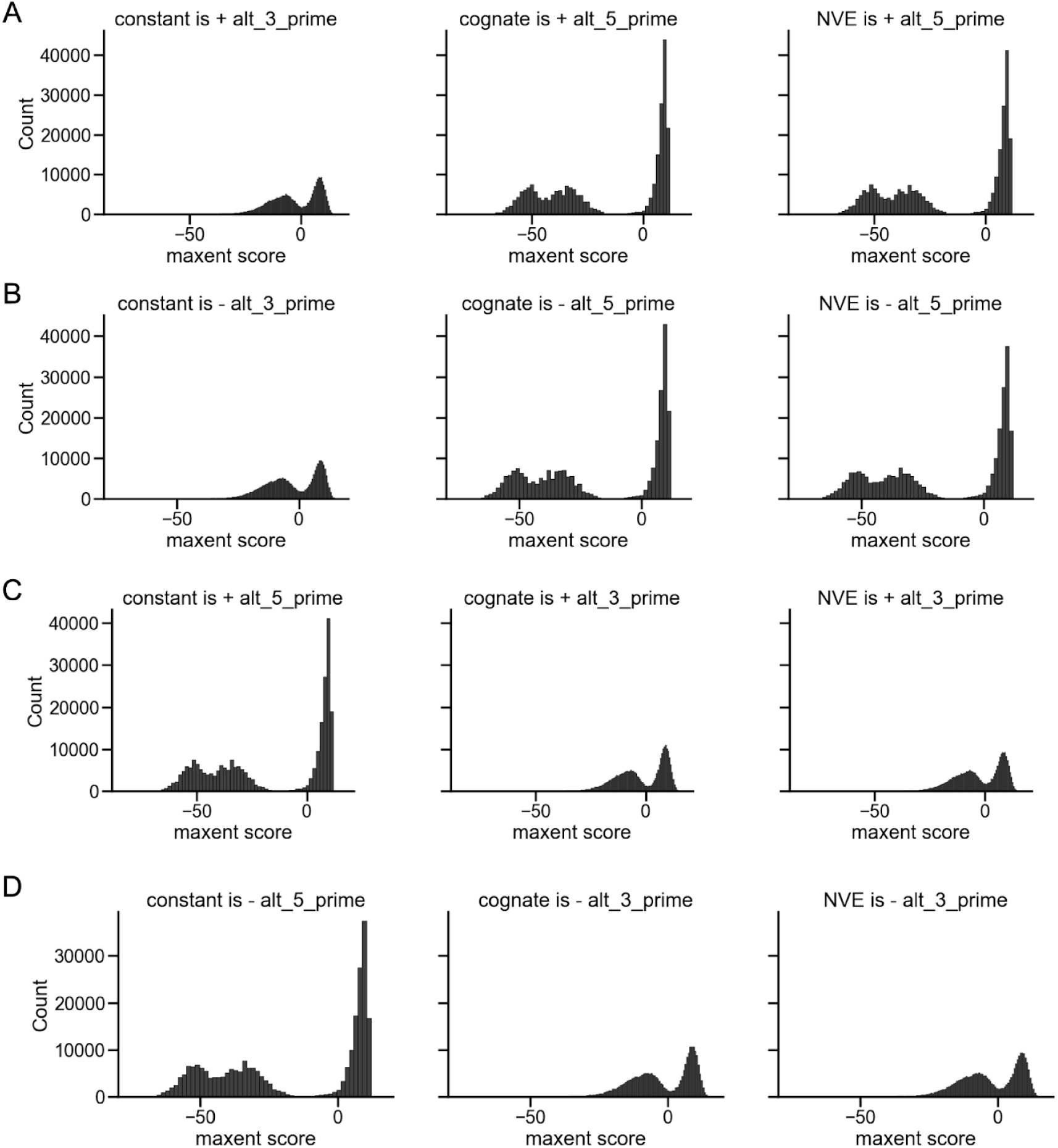
Use of splice site scores to assign type and strand to alternative splice sites. A-D) MaxEnt scores of splice sites, where introns were tested on all possible ways to assign the correct strand and splice site type.

**Supplementary Figure 4.**
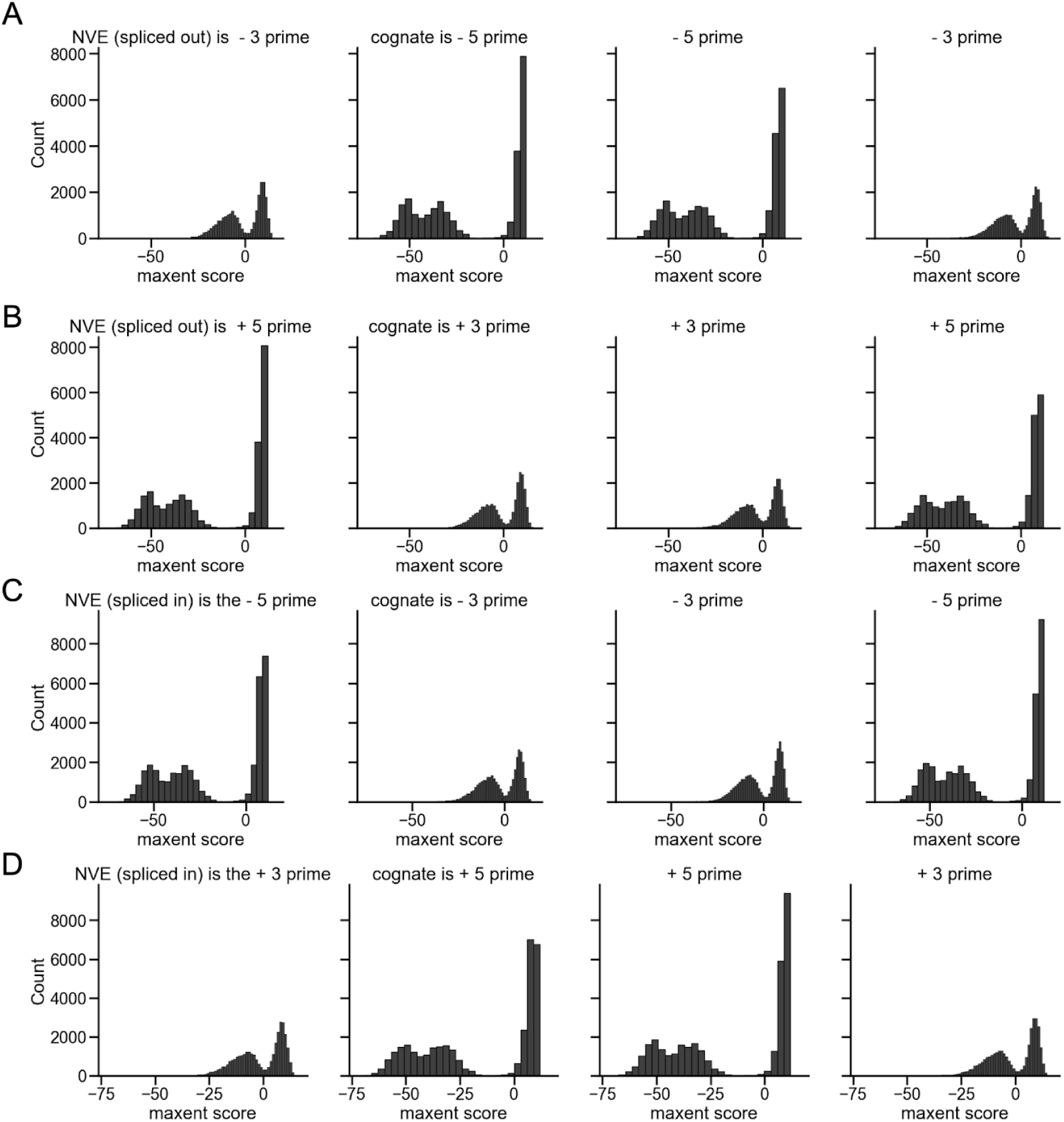
Splice site scores to assign strand to skipped exons. A-D) MaxEnt scores of splice sites, where introns were tested on all possible ways to assign the correct strand and splice site type.

**Supplementary Figure 5.**
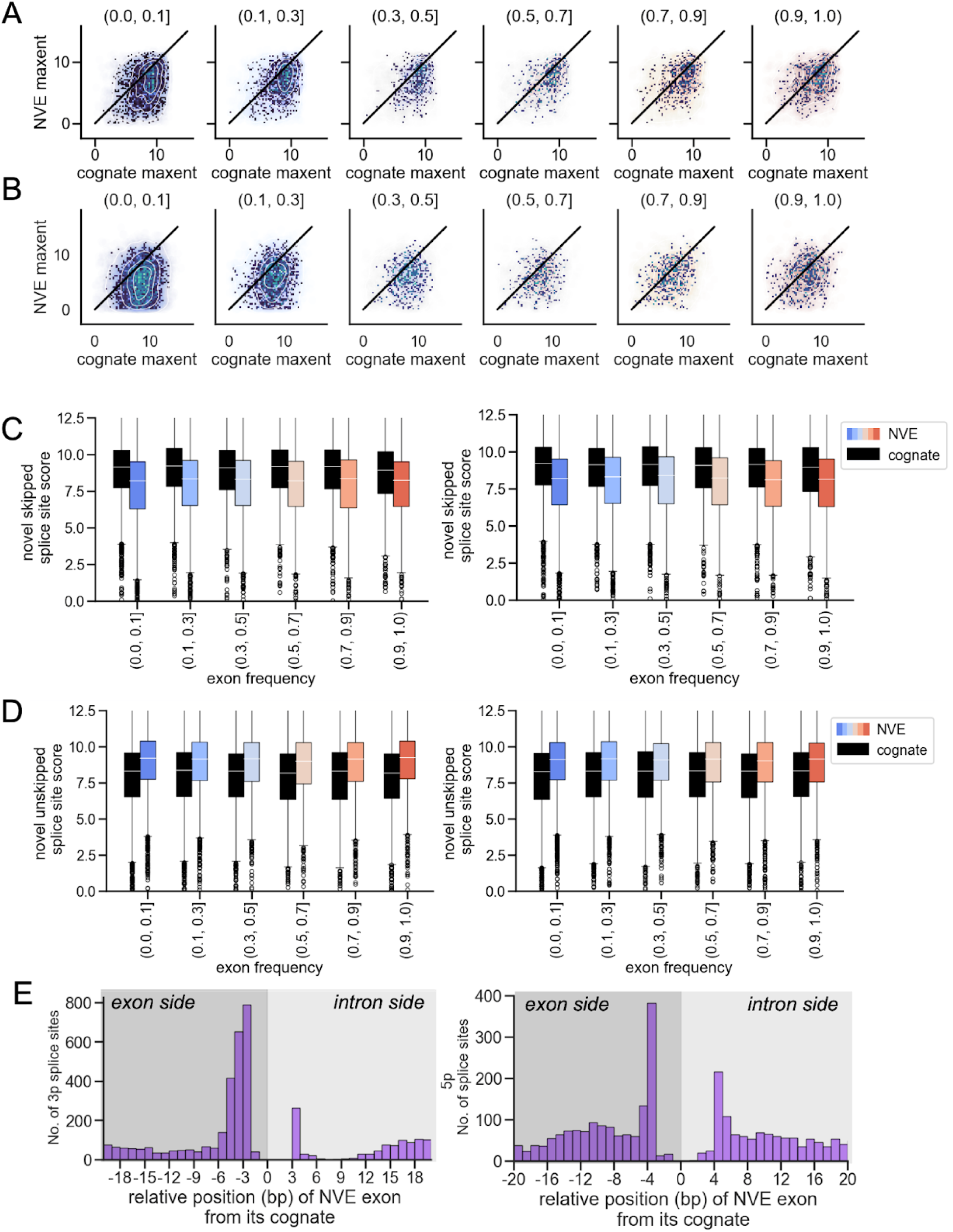
maxent score comparison across EF spectra. A) Correlations between alternative 5’ ss scores between cognate and maxent across EF spectra. Black line represents 1:1 ratio of score. B) Correlations between alternative 3’ ss scores between cognate and maxent across EF spectra. Black line represents 1:1 ratio of score. C-D) Skipped exon scores, stratified on whether an NVE is skipped or unskipped.

## References

1. Wang, E. T. et al. Alternative isoform regulation in human tissue transcriptomes. Nature 456, 470–476 (2008).

2. Martinez, N. M. et al. Alternative splicing networks regulated by signaling in human T cells. RNA 18, 1029–1040 (2012).

3. Irimia, M. et al. A highly conserved program of neuronal microexons is misregulated in autistic brains. Cell 159, 1511–1523 (2014).

4. Hasimbegovic, E. et al. Alternative Splicing in Cardiovascular Disease-A Survey of Recent Findings. Genes 12, (2021).

5. Ren, P. et al. Alternative Splicing: A New Cause and Potential Therapeutic Target in Autoimmune Disease. Front. Immunol. 12, 713540 (2021).

6. Yeo, G. W., Van Nostrand, E., Holste, D., Poggio, T. & Burge, C. B. Identification and analysis of alternative splicing events conserved in human and mouse. Proc. Natl. Acad. Sci. U. S. A. 102, 2850–2855 (2005).

7. Barbosa-Morais, N. L. et al. The evolutionary landscape of alternative splicing in vertebrate species. Science 338, 1587–1593 (2012).

8. Merkin, J. J., Chen, P., Alexis, M. S., Hautaniemi, S. K. & Burge, C. B. Origins and impacts of new mammalian exons. Cell Rep. 10, 1992–2005 (2015).

9. Mazin, P. V., Khaitovich, P., Cardoso-Moreira, M. & Kaessmann, H. Alternative splicing during mammalian organ development. Nat. Genet. 53, 925–934 (2021).

10. Fair, B., et al. Global impact of aberrant splicing on human gene expression levels. bioRxiv (2023) doi:10.1101/2023.09.13.557588.

11. GTEx Consortium. The GTEx Consortium atlas of genetic regulatory effects across human tissues. Science 369, 1318–1330 (2020).

12. Garrido-Martín, D., Borsari, B., Calvo, M., Reverter, F. & Guigó, R. Identification and analysis of splicing quantitative trait loci across multiple tissues in the human genome. Nat. Commun. 12, 727 (2021).

13. Kim Jinkuk et al. Patient-Customized Oligonucleotide Therapy for a Rare Genetic Disease. N. Engl. J. Med. 381, 1644–1652 (2019).

14. Hoffman, G. E. et al. CommonMind Consortium provides transcriptomic and epigenomic data for Schizophrenia and Bipolar Disorder. Sci Data 6, 180 (2019).

15. Wilks, C. et al. recount3: summaries and queries for large-scale RNA-seq expression and splicing. Genome Biol. 22, 323 (2021).

16. Jaganathan, K. et al. Predicting Splicing from Primary Sequence with Deep Learning. Cell 176, 535–548.e24 (2019).

17. Wagner, N. et al. Aberrant splicing prediction across human tissues. Nat. Genet. 55, 861–870 (2023).

18. Dawes, R. et al. SpliceVault predicts the precise nature of variant-associated mis-splicing. Nat. Genet. 55, 324–332 (2023).

19. Bénitière, F., Necsulea, A. & Duret, L. Random genetic drift sets an upper limit on mRNA splicing accuracy in metazoans. Elife 13, (2024).

20. Schafer, S. et al. Alternative Splicing Signatures in RNA-seq Data: Percent Spliced in (PSI). Curr. Protoc. Hum. Genet. 87, 11.16.1–11.16.14 (2015).

21. Baek, D. & Green, P. Sequence conservation, relative isoform frequencies, and nonsense-mediated decay in evolutionarily conserved alternative splicing. Proc. Natl. Acad. Sci. U. S. A. 102, 12813–12818 (2005).

22. Yeo, G. & Burge, C. B. Maximum entropy modeling of short sequence motifs with applications to RNA splicing signals. J. Comput. Biol. 11, 377–394 (2004).

23. Karczewski, K. J. et al. The mutational constraint spectrum quantified from variation in 141,456 humans. Nature 581, 434–443 (2020).

24. Kanai, M. et al. Insights from complex trait fine-mapping across diverse populations. bioRxiv (2021) doi:10.1101/2021.09.03.21262975.

25. Lek, M. et al. Analysis of protein-coding genetic variation in 60,706 humans. Nature 536, 285–291 (2016).

26. Nioi, P. et al. Variant ASGR1 Associated with a Reduced Risk of Coronary Artery Disease. N. Engl. J. Med. 374, 2131–2141 (2016).

27. Ali, L. et al. Common gene variants in ASGR1 gene locus associate with reduced cardiovascular risk in absence of pleiotropic effects. Atherosclerosis 306, 15–21 (2020).

28. Kolakada, D. et al. A system of reporters for comparative investigation of EJC-independent and EJC-enhanced nonsense-mediated mRNA decay. Nucleic Acids Res. 52, e34 (2024).

29. Fiszbein, A., Krick, K. S., Begg, B. E. & Burge, C. B. Exon-Mediated Activation of Transcription Starts. Cell 179, 1551–1565.e17 (2019).

30. Botto, A. E. C. et al. Reciprocal regulation between alternative splicing and the DNA damage response. Genet. Mol. Biol. 43, e20190111 (2020).

31. García-Pérez, R. et al. The landscape of expression and alternative splicing variation across human traits. Cell Genom 3, 100244 (2023).

32. Mariotti, M., Kerepesi, C., Oliveros, W., Mele, M. & Gladyshev, V. N. Deterioration of the human transcriptome with age due to increasing intron retention and spurious splicing. bioRxiv 2022.03.14.484341 (2022) doi:10.1101/2022.03.14.484341.

33. Bhadra, M., Howell, P., Dutta, S., Heintz, C. & Mair, W. B. Alternative splicing in aging and longevity. Hum. Genet. 139, 357–369 (2020).

34. Lai, R. W. et al. Multi-level remodeling of transcriptional landscapes in aging and longevity. BMB Rep. 52, 86–108 (2019).

35. Li, H., Wang, Z., Ma, T., Wei, G. & Ni, T. Alternative splicing in aging and age-related diseases. Translational Medicine of Aging 1, 32–40 (2017).

36. Hartmann, C. et al. Systematic estimation of biological age of in vitro cell culture systems by an age-associated marker panel. Front Aging 4, 1129107 (2023).

37. Wang, Q. S. et al. Leveraging supervised learning for functionally informed fine-mapping of cis-eQTLs identifies an additional 20,913 putative causal eQTLs. Nat. Commun. 12, 3394 (2021).

38. Barbeira, A. N. et al. Exploiting the GTEx resources to decipher the mechanisms at GWAS loci. Genome Biol. 22, 1–24 (2021).

39. Fairbrother, W. G. et al. RESCUE-ESE identifies candidate exonic splicing enhancers in vertebrate exons. Nucleic Acids Res. 32, W187–90 (2004).

40. Wang, Z. et al. Systematic identification and analysis of exonic splicing silencers. Cell 119, 831–845 (2004).

41. ENCODE Project Consortium et al. Expanded encyclopaedias of DNA elements in the human and mouse genomes. Nature 583, 699–710 (2020).

42. Van Nostrand, E. L. et al. A large-scale binding and functional map of human RNA-binding proteins. Nature 583, 711–719 (2020).

43. Westra, H.-J. & Franke, L. From genome to function by studying eQTLs. Biochim. Biophys. Acta 1842, 1896–1902 (2014).

44. Wuttke, M. et al. A catalog of genetic loci associated with kidney function from analyses of a million individuals. Nat. Genet. 51, 957–972 (2019).

45. Trubetskoy, V. et al. Mapping genomic loci implicates genes and synaptic biology in schizophrenia. Nature 604, 502–508 (2022).

46. Teumer, A. et al. Genome-wide association meta-analyses and fine-mapping elucidate pathways influencing albuminuria. Nat. Commun. 10, 4130 (2019).

47. Xu, Y.-X. et al. EDEM3 Modulates Plasma Triglyceride Level through Its Regulation of LRP1 Expression. iScience 23, 100973 (2020).

48. Neil, E. E. & Bisaccia, E. K. Nusinersen: A Novel Antisense Oligonucleotide for the Treatment of Spinal Muscular Atrophy. J. Pediatr. Pharmacol. Ther. 24, 194–203 (2019).

49. Lim, K. H. et al. Antisense oligonucleotide modulation of non-productive alternative splicing upregulates gene expression. Nat. Commun. 11, 3501 (2020).

50. Walker, R. L. et al. Genetic Control of Expression and Splicing in Developing Human Brain Informs Disease Mechanisms. Cell 179, 750–771.e22 (2019).

51. Trabzuni, D. et al. Widespread sex differences in gene expression and splicing in the adult human brain. Nat. Commun. 4, 2771 (2013).

52. Blekhman, R., Marioni, J. C., Zumbo, P., Stephens, M. & Gilad, Y. Sex-specific and lineage-specific alternative splicing in primates. Genome Res. 20, 180–189 (2010).

53. Scotti, M. M. & Swanson, M. S. RNA mis-splicing in disease. Nat. Rev. Genet. 17, 19–32 (2016).

54. Li, Y. I. et al. Annotation-free quantification of RNA splicing using LeafCutter. Nat. Genet. 50, 151–158 (2018).

55. Carpenter, B. Hierarchical partial pooling for repeated binary trials. https://mc-stan.org/users/documentation/case-studies/pool-binary-trials.html (2016).

56. Young-Xu, Y. & Chan, K. A. Pooling overdispersed binomial data to estimate event rate. BMC Med. Res. Methodol. 8, 58 (2008).

57. Wainberg, M., Alipanahi, B. & Frey, B. Does conservation account for splicing patterns? BMC Genomics 17, 787 (2016).

58. Benoit Bouvrette, L. P., Bovaird, S., Blanchette, M. & Lécuyer E. oRNAment: a database of putative RNA binding protein target sites in the transcriptomes of model species. Nucleic Acids Res. 48, D166–D173 (2020).

